# RAV1 mediates cytokinin signalling for regulating primary root growth in Arabidopsis

**DOI:** 10.1101/2022.07.13.499994

**Authors:** Drishti Mandal, Saptarshi Datta, Giridhar Ravindra, Pranab Kumar Mondal, Ronita Nag Chaudhuri

## Abstract

Root growth dynamics is an outcome of complex hormonal crosstalk. The primary root meristem size for example, is determined by antagonizing actions of cytokinin and auxin. Here we show that RAV1, a member of the AP2/ERF family of transcription factors, mediates cytokinin signalling in roots to regulate meristem size. The *rav1* mutants have prominently longer primary roots, with a meristem that is significantly enlarged and contain higher cell numbers, compared to wild type. The mutant phenotype could be restored on exogenous cytokinin application or by inhibiting auxin transport. At the transcript level, primary cytokinin-responsive genes like *ARR1, ARR12* were significantly downregulated in the mutant root, indicating impaired cytokinin signalling. In concurrence, cytokinin induced regulation of *SHY2*, an Aux/IAA gene, and auxin efflux carrier *PIN1* was hindered in *rav1*, leading to altered auxin transport and distribution. This effectively altered root meristem size in the mutant. Notably, CRF1 another member of the AP2/ERF family implicated in cytokinin signalling, is transcriptionally repressed by RAV1 to promote cytokinin response in roots. Further correlating RAV1 to cytokinin signalling, our results demonstrate that cytokinin upregulate *RAV1* expression through ARR1, during post-embryonic root development. Regulation of *RAV1* expression is a part of secondary cytokinin response that eventually represses *CRF1* to augment cytokinin signalling. To conclude, in Arabidopsis, RAV1 functions in a branch pathway downstream to ARR1 that regulates *CRF1* expression to enhance cytokinin action during primary root development.

## Introduction

The root architecture of a plant critically influences its growth and adaptability under various developmental and environmental cues. Morphology of the root structure itself is very consistent in a particular species. However, the spatial configuration of the roots, i.e., number, position and growth direction of primary, lateral and adventitious roots (wherever applicable) which comprises the root system architecture (RSA) is highly variable (Giehl and von Wirén, 2014). The basic structure remaining same, root architecture undergoes continuous manipulation based on ever-changing environmental and growth parameters (Malamy, 2005; Giehl and von Wirén, 2014; Tardieu et al., 2017; Perianez-Rodriguez et al., 2021). RSA evolves during post-embryonic growth and the cues perceived over time are gradually integrated into an intrinsic root development program (Rosas et al., 2013; Kellermeier et al., 2014; Pandey et al., 2021). This adaptive dynamism in root growth is the key to proper health and vigour of the above-ground biomass (Paez-Garcia et al., 2015; Rogers and Benfey, 2015; Lombardi et al., 2021) and warrants our thorough understanding.

Like any other developmental process, root development too is dependent on an intricate functional network of several hormone signalling pathways (Petricka et al., 2012; Liu et al., 2014; Pacifici et al., 2015; Wachsman et al., 2015). In Arabidopsis, the primary root can be discretely distinguished into four zones such as meristematic zone, transition zone, elongation zone and maturation zone (Verbelen et al., 2006). Each zone is a consequence of the interplay of a specific set of hormones (Ubeda-Tomás et al., 2012; Jung and McCouch, 2013; Lee et al., 2013; Qin et al., 2019; Zluhan-Martínez et al., 2021). A critically coordinated action between auxin and cytokinin signalling for example, forms the basis of cell proliferation regulation and differentiation during primary root growth. Polar transport of auxin and generation of correct auxin gradient is essential for cell division in root meristem, while subsequent cell differentiation is promoted by cytokinin (Dello Ioio et al., 2007). Cytokinin is known to act at the transition zone to limit auxin transport and action primarily to the root meristematic zone through regulation of SHY2, a negative regulator of auxin signalling (Tian et al., 2002; Moubayidin et al., 2009; Ruzicka et al., 2009; Moubayidin et al., 2010). In Arabidopsis roots, SHY2 acts to reduce expression of the *PIN* genes that function as auxin efflux carrier and consequently helps to maintain precise auxin gradient for proper root patterning (Blilou et al., 2005; Dello Ioio et al., 2007; Dello Ioio et al., 2008; Dello Ioio et al., 2008; Šimášková et al., 2015; Li et al., 2020). Consequently, mutations in cytokinin biosynthesis genes or signalling molecules show enlarged root meristem and longer primary root growth (Moubayidin et al., 2010). This dual antagonism between auxin and cytokinin at the junction of meristematic zone and elongation zone, is the key factor that decides primary root length in Arabidopsis - a critical parameter that influences plant health and survivability under control as well as non-congenial conditions.

RAV1, a member of the AP2/ERF as well as B3 superfamily is unique in having two DNA binding domains, making it one of the rarest kinds of transcription factor in the plant family, till date. It has an N-terminal AP2 domain and a C-terminal B3 domain, by virtue of which it has the ability to bind to CAACA and CACCTG cis elements, respectively (Kagaya et al., 1999; Feng et al., 2005). Scattered evidences have indicated RAV1 to be involved in certain developmental and stress responsive pathways (Hu et al., 2004; Sohn et al., 2006; Kagaya and Hattori, 2009; Woo et al., 2010; Feng et al., 2014; Fu et al., 2014; Shin and Nam, 2018; Ren et al., 2021). To unravel the role of RAV1 in regulating dehydration stress response in Arabidopsis, it has been shown before that absence of RAV1 leads to enhanced primary root growth (Sengupta et al., 2020). Consequently, *rav1* mutant plants had reduced water loss and showed more efficient response to dehydration stress, compared to wild type. The present work addresses the mechanism of RAV1 mediated regulation of primary root growth. Our findings suggest that during post-embryonic root development, RAV1 augments cytokinin signalling for controlling primary root length in Arabidopsis.

## Results

### Absence of *RAV1* causes increase in root meristem size

It has been earlier shown that RAV1 deletion mutants have longer primary root growth compared to wild type, within 14 days after germination (dag) (Sengupta et al., 2020). To understand the basis of such differential root growth, we monitored primary root length in RAV1 deletion mutants in comparison to wild type from 1 to 14 dag. For this we worked with two mutants of RAV1 named hereafter as *rav1* (N420832) and *rav1-1* (N655012). Both the mutants have T-DNA insertion within the single exon that encodes for RAV1 in Arabidopsis (Supplemental Figure S1A). As reported previously (Sengupta et al., 2020), it was observed that the primary root growth was higher in both the RAV1 deletion mutants compared to wild type (Figure 1A, Supplemental Figure S1B, S1C). The *rav1* roots grew longer than wild type starting from 4 dag, and the difference in root length became more significant with increased time after germination (Figure 1B). While both *rav1* and *rav1-1* exhibited longer primary root compared to wild type, *rav1-1* seeds showed more delayed germination and delayed transition from vegetative to reproductive phase, compared to *rav1*. Hence, keeping future application prospects in mind, all the subsequent experiments performed hereafter were done using *rav1*. Next, we probed at the cellular level, to understand which region(s) of the primary root in *rav1* was different from wild type. For this we performed confocal microscopy with roots of *rav1* and wild type, as mentioned in the Materials and Methods section. Our observations revealed that the root meristem size of the mutants was larger compared to wild type (Figure 1C). Root meristem size is expressed as the number of cells in cortex files that lies between the quiescent centre (QC) and the first elongated cell (Dello Ioio et al., 2007). Results indicated that in wild type primary roots while the meristem size remained limited to about 34 (± 1.5) cells within 5-7 dag, in *rav1* roots the cell number was more than 40 (± 2.5) within 5 dag and increased further around 7 dag (Figure 1D). It was thus evident that in wild type primary root meristem size gets ascertained and restricted within around 5 dag. On the contrary, in absence of RAV1 the primary root meristem continues to grow even 7 dag. Collectively, the above results indicate that absence of RAV1 results in longer primary root growth due to increased root meristem size, compared to wild type.

**Figure 1.**
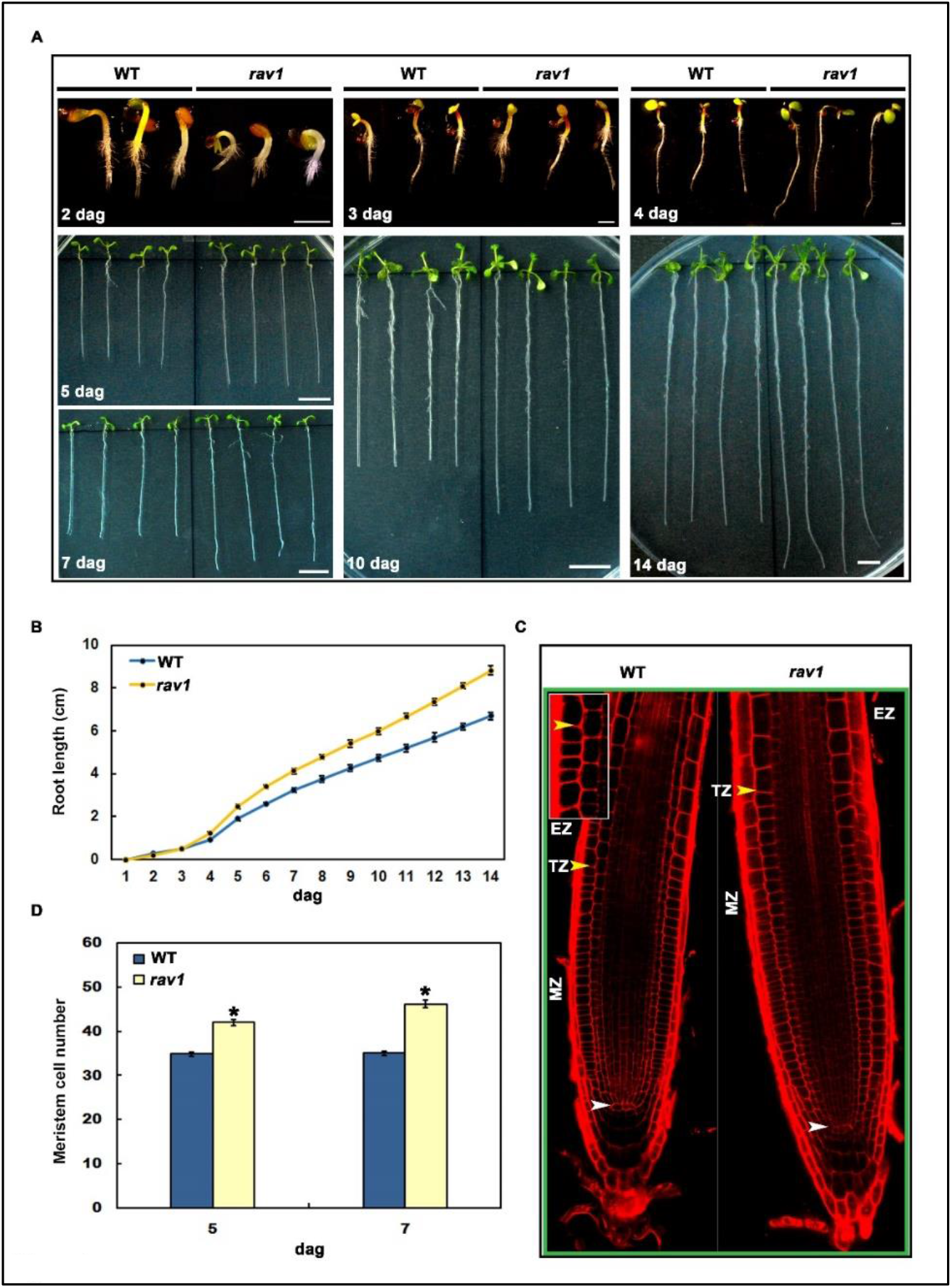
The *rav1* mutant exhibits a longer root length phenotype. **A**. Primary root growth of wild-type (WT) and *rav1* grown vertically on MS medium for 2, 3, 4, 5, 7, 10 and 14 days after germination (dag). Scale bars for 2-4 dag = 1 mm and for 5-14 dag = 1cm. **B**. Primary root length measurements of WT and *rav1* mutants over time. The data represent the mean of three replicate experiments with n ≥ 30 seedlings for each replicate of each genotype with standard error of mean (SE). **C**. Longitudinal view of propidium iodide-stained 7-day old WT and *rav1* root meristem, where white and yellow arrowheads indicate, the quiescent center (QC) and the cortex transition zone (TZ), respectively; MZ = meristematic zone and EZ= elongation zone. The transition zone in WT root cortex is shown in the inset. **D**. Root meristem cell number of WT and *rav1* plants 5 and 7 days after germination (dag). Cortical meristematic cells enclosed between the white and yellow arrowheads (C) were counted and represented as meristem cell number. Results represent means from at least three independent replicates (n ≥ 20) and error bars represent standard error of mean (SE). Student’s t-test with paired two-tailed distribution was used for statistical analysis and P ≤ 0.05 was denoted by *.

### *rav1* mutant phenotype can be reverted by exogenous cytokinin treatment

The hormone cytokinin has been implicated in regulating primary root meristem size through strict regulation of auxin action and control over cell differentiation (Dello Ioio et al., 2007; Dello Ioio et al., 2008; Dello Ioio et al., 2008; Kuderová et al., 2008; Ruzicka et al., 2009; Moubayidin et al., 2010; Šimášková et al., 2015). We therefore next examined, whether cytokinin has a role to play in the differential root growth observed between *rav1* and wild type seedlings. For this, wild type and *rav1* seeds were germinated in presence of different concentrations of exogenous cytokinin (6-Benzylaminopurine, 6-BAP). As shown in Figure 2A, it is apparent that when germinated in presence of 6-BAP, both *rav1* and wild type seedlings showed reduction in primary root growth. Thus, exogenous cytokinin treatment could reduce primary root growth in both *rav1* and wild type. For comparison, we also checked the primary root length of cytokinin response mutant *arr1* under similar growth conditions and observed reduction in primary root length in presence of 6-BAP (Figure 2A). This is in consonance with previous reports that have shown that treatment with exogenous cytokinin reduces root length and meristem size even in cytokinin signalling mutants (Sakai et al., 2001; Raines et al., 2016). However, it is to be noted that under same concentrations of 6-BAP reduction in primary root length of wild type was greater, compared to both *rav1* and *arr1* (Figure 2B). This implies that compared to wild type, both *arr1* and *rav1* roots are less sensitive to cytokinin. Microscopic analyses with 6-BAP treated seedlings further confirmed that upon exogenous cytokinin application root meristem size in both *rav1* and wild type seedlings became shorter, compared to when germinated under control conditions (Figure 2C). Again, the reduction in meristem size was higher in wild type roots compared to *rav1*, in presence of same concentration of exogenous cytokinin. Moreover, in presence of 10 nM 6-BAP meristem cell numbers of *rav1* primary roots was found to be comparable to roots of wild type seedlings grown under control conditions (Figure 2D). The results above thus indicate that enhanced root length and meristem size observed in *rav1* is due to impaired cytokinin response.

**Figure 2.**
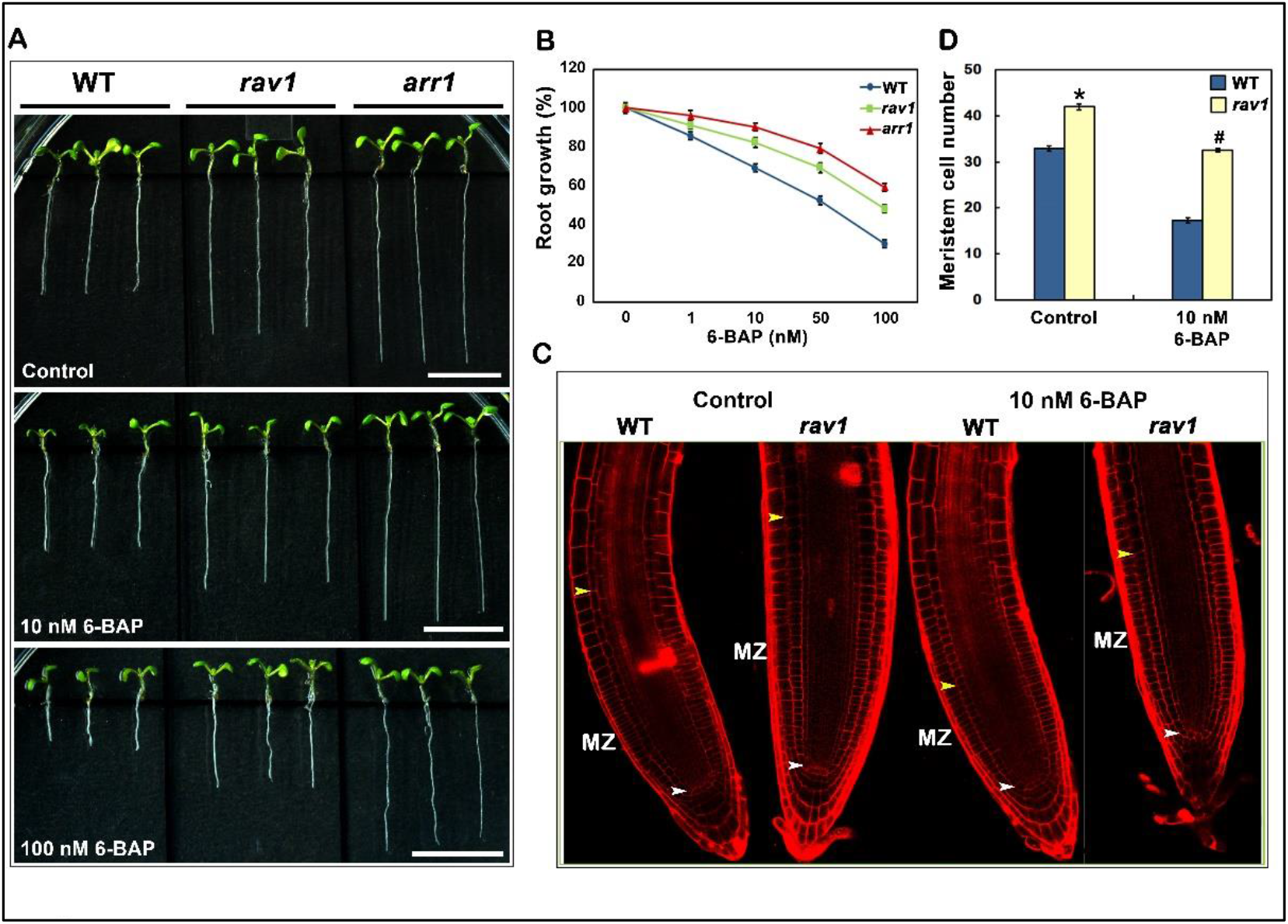
Effect of exogenous cytokinin on primary root growth. **A**. Morphology of wild type, *rav1* and *arr1* seedlings 5 dag in presence of either 6-Benzylaminopurine (6-BAP) (concentration 10 nM or 100 nM) or 0.1 % DMSO (control). Scale bars = 1 cm. **B**. Root growth (%) of WT, *rav1* and *arr1* mutants treated with different concentrations of 6-BAP, relative to control. The data presented is an average of three biological replicates, with each replicate having n ≥ 30 seedlings and error bars representing standard error of mean (SE). **C**. Longitudinal view of root meristem of WT and *rav1* seedlings 5 dag, grown in presence or absence of 10 nM 6-BAP; white and yellow arrowheads indicate, QC and TZ respectively, MZ = Meristematic zone. **D**. Root meristem cell number of WT and *rav1* seedlings 5 dag. Cortex meristematic cells enclosed between the white and yellow arrowheads (C) were counted and represented as meristem cell number. Data represent means from three independent replicates (n ≥ 20) and error bars represent SE. Anova two-factor with replication method was used to measure the variance and the results with P value ≤ 0.05 considered to be significant. Variance between WT and *rav1* are denoted by * while variance within *rav1* samples is denoted by **#**.

### Cytokinin application specifically to the *rav1* root alters root meristem size

By convention, studies that involve rescue of phenotypes resulting from defects in hormone signalling or biosynthesis, allow seed germination or treatment of the entire plant or plant part in presence of specific hormones or hormone inhibitor. It is known that phytohormones like auxin or cytokinin can play diverse roles in different plant parts or different zones of the same organ, during various developmental stages. Hence germinating seeds in presence of a particular hormone, or treatment of the entire plant may overshadow the precise effects expected to be endowed by the hormone on development of a specific plant part, like root. For a complex and dynamic organ like primary root, development of different zones is known to be an outcome of interplay between specific sets of hormones (Verbelen et al., 2006; Dello Ioio et al., 2007; Dello Ioio et al., 2008; Dello Ioio et al., 2008; Ruzicka et al., 2009; Moubayidin et al., 2010; Šimášková et al., 2015). To address this issue pertaining to our study, we used a microfluidic platform, referred to as the plant root microfluidic system (PRMS) hereafter, to treat specific region of *rav1* roots with cytokinin. For this, wild type and *rav1* plants were grown in PRMS, as described in the Materials and methods section (Figure 3A, Supplemental Figure S2A). We first compared wild type and mutant plant root growth under similar conditions on agar plates and in the microfluidics system. As reported earlier (Meier et al., 2010), growth rate of both wild type and mutant seedlings was reduced by more than 50% in the confined space of microfluidic channel, compared to that on agar plates (Supplemental Figure S2B, S2C, S2D). Despite such reduction in growth rate in the PRMS, primary root length of *rav1* was significantly higher, compared to wild type, as observed earlier on agar plates (Supplemental Figure S2D). After monitoring growth under control conditions, next attempt taken was to check the effect of cytokinin treatment on *rav1* roots in the microfluidic channel. As effect of cytokinin on root meristem and differentiation zone occurs primarily within 2/3-4/5 days after germination, we transferred wild type and *rav1* seedlings to the PRMS 3 dag. The plants were allowed to grow therein for next 12 hours. After 12 hours, 5 μM 6-BAP supplemented MS media was supplied through the microfluidic channel to WT and *rav1* roots. 6-BAP treatment was done for 1 to 5 hr, such that the hormone reached specifically to a region of the root that is within 1 mm from the root tip, encompassing primarily the meristematic, transition and elongation zones (Verbelen et al., 2006) (Figure 3A & 3B). Interestingly, results indicated that for *rav1* roots that grew in presence of 5 μM 6-BAP for 5 hr, the length of root meristem was reduced significantly, compared to *rav1* that grew for the same time period under control conditions (Figure 3B & 3C). Furthermore, the length of 5 hr 6-BAP-treated *rav1* root meristem was similar to that of wild type grown under control conditions (Figure 3C). As depicted in Figure 3D, this data implied that the increased root growth and meristem size observed in *rav1* root could be arrested upon exogenous cytokinin treatment. To conclude, the plant root microfluidic system (PRMS) allowed us to precisely treat specific regions of the *rav1* root with cytokinin for defined time periods and to monitor its effect on the root length and meristem size. We could demonstrate that few hours of cytokinin treatment to *rav1*, 3 days post germination, could restrict the root growth and the meristematic zone length in the mutant, such that it became comparable to that of untreated wild type roots. From the above results it becomes further evident that in absence of RAV1, cytokinin action in primary root is impaired such that proper regulation on root meristem size is effectively lost.

**Figure 3.**
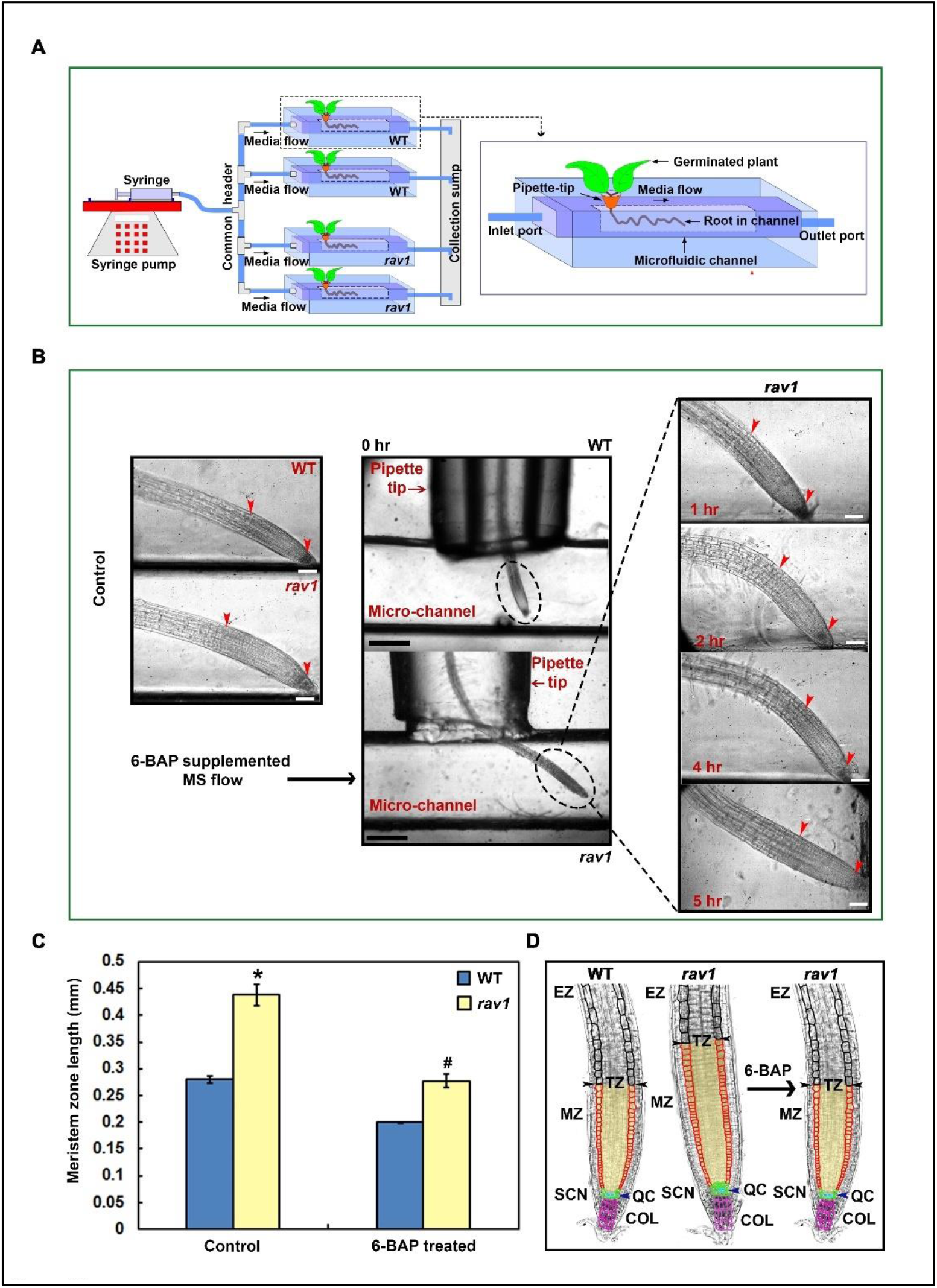
Exogenous cytokinin application to *rav1* roots in a Plant Root Microfluidic System. **A**. Schematic diagram showing the plant root microfluidic system (PRMS). Four microfluidic channels in PRMS were connected to a common header through the inlet ports and the outlet ports were extended to the collection sump using connectors. This setup is connected to syringe pump for supplying media to the microfluidic channels. Right panel showing an enlarged view of a microfluidic channel with germinating plant and root growing into the channel. **B**. Microfluidics channels with WT and *rav1* roots. Surface sterilized Arabidopsis seeds were germinated in 5 mm long pipette tip cone embedded in solid MS medium (3 % sucrose). 3 dag the pipette tip cone containing seedlings were inserted into microfluidic channel and supplied with ½-strength MS media for 12 hours. WT and *rav1* roots were thereafter supplied with ½-strength MS media supplemented with 6-BAP (5 μM) for 5 hr. WT and *rav1* growing in presence of ½-strength MS media supplemented with 0.1 % DMSO for 5 hr was used as control. Left and middle panels show 20X and 5X images respectively, of roots of WT and *rav1* seeds grown under control conditions. Right panel shows roots of *rav1* mutants after 1, 2, 4, and 5 hr of 6-BAP treatment. Red arrows indicate meristematic zone. Scale bar left and right panel = 0.1 mm, middle panel = 0.5 mm. **C**. Meristematic zone length measurements of WT and *rav1* plants treated with 6-BAP for 5 hr. Data represent means from three independent replicates and error bars represent SE. Anova two-factor with replication method was used to measure the variance and the results with P value ≤ 0.05 were considered to be significant. Variance in between WT and *rav1* is denoted by * and variance in between *rav1* samples is denoted by **#. D**. Schematic representation of longitudinal view of a part of Arabidopsis root showing exogenous cytokinin (6-BAP) application to the *rav1* roots altered root meristem size. The stem cell niche (SCN), quiescent center (QC), columella cells (COL) are marked in green, blue and pink, respectively; a cortex cell file of the meristematic zone and elongation zone (EZ) are marked in red and black, respectively. Black arrows indicate the transition zone (TZ). Meristematic zone is denoted as MZ.

### Auxin transport and distribution is altered in the roots of *rav1* plants

Previous works have already demonstrated that root meristem size is negatively regulated by cytokinin (Dello Ioio et al., 2007) and that it acts through regulation of polar auxin transport (Ruzicka et al., 2009). As *rav1* roots have impaired cytokinin response, we next wanted to check whether auxin distribution or transport was altered in the *rav1* roots. For this, auxin responsive promoter DR5::GFP construct was introduced into wild type and *rav1* roots through hairy root transformation and GFP expression was monitored. We observed that DR5 expression in *rav1* roots that were transformed 2 dag, showed enhanced auxin response, compared to wild type (Figure 4A, upper panel). Furthermore, in wild type roots that were transformed 7 dag, the DR5 expression primarily remained restricted to the meristematic zone, while in *rav1* roots the expression was distributed towards the elongation zone (Figure 4A, lower panel), again indicating increased auxin response. It is to be noted that GFP expression in pCAMBIA1304 vector under 35S promoter, showed no difference in wild type and *rav1* roots (Supplemental Figure S3). This confirmed that the difference observed with DR5 expression is due to differential auxin response in *rav1* roots, compared to wild type. It is thus evident that auxin distribution and response is differentially regulated in *rav1* roots, compared to wild type. In consonance with this result, we further found that proPIN1::GFP expression in the *rav1* and wild type roots was differential. In wild type roots the proPIN1::GFP expression was primarily enriched in the root tip and in the vascular tissue. In *rav1* roots however, the proPIN1::GFP expression spanned much higher up from the root tip and had a more extensive distribution towards the elongation zone, compared to wild type (Figure 4B). One of the most widely accepted fact is that, in roots cytokinin controls auxin transport through transcriptional and/or post-transcriptional regulation of *PIN1*, the auxin efflux carrier (Dello Ioio et al., 2008; Zhang et al., 2011). PIN1 works towards maintaining right auxin concentration in specific regions of the root, through regulation of directional flow of auxin (Friml et al., 2002; Vieten et al., 2005; Benjamins and Scheres, 2008). Hence, in the *rav1* roots altered *PIN1* expression has generated an altered auxin response. Thus, impaired cytokinin response in *rav1* roots, through altered PIN1 expression has led to differential auxin gradient in *rav1* roots, compared to wild type. Such enhanced auxin distribution and response in *rav1* roots consequently generated a larger root meristem relative to wild type.

**Figure 4.**
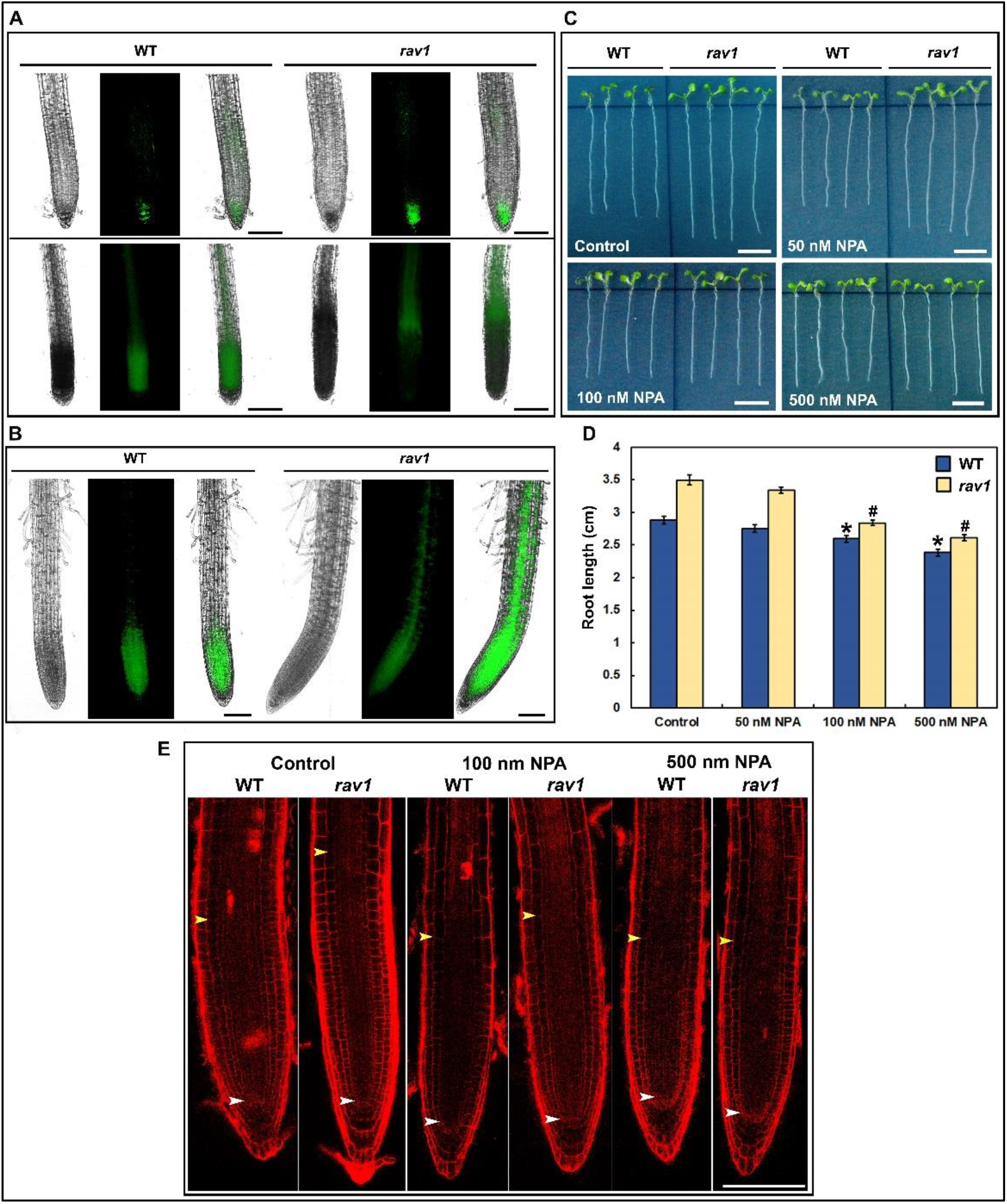
Auxin transport and distribution in the roots of *rav1* plants. **A**. Localization of DR5::GFP expression in wild type and *rav1* roots using hairy root transformation, 2 dag (Upper panel) and 7 dag (lower panel). seedlings were used for hairy root transformation. Scale bars = 100 µm. **B**. Localization of PIN1::GFP expression in wild type and *rav1* roots using hairy root transformation in 7 dag seedling. Scale bars = 100 µm. **C**. Wild type and *rav1* root growth, in presence of either *N*-1-naphthylphthalamic acid (NPA) (concentration 50 nM, 100 nM or 500 nM) or 0.1 % DMSO (control), 6 dag. Scale bars = 1 cm. **D**. Root length in wild type and *rav1* mutants in absence or presence of NPA. The data presented is an average of three biological replicates, with each replicate having n ≥ 20 seedlings and error bars representing standard error (SE). Student’s t-test with paired two-tailed distribution was used for statistical analysis and P ≤ 0.05 was marked. Variance within WT and *rav1* samples are denoted by * and #, respectively. **E**. Longitudinal view of root meristem of WT and *rav1* seedlings grown in presence or absence of NPA (100 nM or 500 nM), white and yellow arrowheads indicate QC and TZ respectively.

*N*-1-naphthylphthalamic acid (NPA) is known to be an auxin transport inhibitor that specifically inhibits PIN1 action (Fujita and Syono, 1996; Yuan and Huang, 2016; Abas et al., 2021; Teale et al., 2021). We therefore argued that, if *rav1* roots were longer due to inefficient cytokinin action and consequently altered auxin transport, then NPA treatment of *rav1* seedlings should affect its primary root length. Therefore, wild type and *rav1* seeds were germinated in presence of different concentrations of NPA and root lengths were measured 6 dag (Figure 4C & 4D). Results indicated that when germinated in presence of NPA, primary root length of both wild type and *rav1* was reduced. It is implicative from this data that NPA could inhibit PIN1 action and consequently root growth in both wild type and *rav1* seedlings. However, *rav1* roots with more extensive auxin distribution showed more significant root length reduction compared to wild type. Confocal microscopic analyses further revealed that germination in presence NPA reduced meristem zone length in *rav1* roots (Figure 4E). Markedly, in presence of 100 nM and 500 nM NPA, root meristem length reduction in *rav1* was significantly higher compared to wild type. We have noted earlier (Figure 1), that in wild type seedlings the root meristem size reaches a plateau around 5 dag, while in *rav1* it continues to grow thereafter. Thus, it seems logical that effect of NPA was more significant on *rav1* root length and meristem size, compared to wild type, when monitored 6 dag. From all the above data, it can thus be summed up that *rav1* primary roots were longer than that of wild type with a larger meristem size due to alteration in auxin transport and distribution.

### Expression of cytokinin-responsive genes is altered in *rav1* roots

To gain further insight into the molecular mechanism of RAV1 mediated control of primary root length, gene expression analyses were done hereafter. For this, RNA was isolated from the roots of *rav1* and wild type seedlings, 3-14 dag, and qRT-PCR was performed. As seen from Figure 5A, *RAV1* gene expression reduced significantly by 5 dag relative to 3 dag, increasing further again around 14 dag. This indicated that the *RAV1* gene has a specific expression pattern during different stages of root development following germination. Subsequently, to understand why cytokinin action was inefficient in *rav1* roots, we thought that it was essential to know whether it was cytokinin biosynthesis or cytokinin signalling that was hindered in absence of RAV1. Therefore, we checked important marker genes for cytokinin biosynthesis such as *IPT3, IPT5* and *IPT7* (Miyawaki et al., 2004; Dello Ioio et al., 2007; Ivanov and Filin, 2018) and for cytokinin signalling like *ARR1, ARR12* and *ARR6*. It has been previously reported that *ARR1* and *ARR12* are type-B ARR genes that get expressed mainly in the transition zone of primary roots at the early stages of development, after seed germination (Dello Ioio et al., 2007). *ARR6*, a type-A ARR gene, is one of the immediate early -response cytokinin gene which is a direct target of ARR1 and other type-B ARRs (Hwang and Sheen, 2001; Sakai et al., 2001; Rashotte et al., 2003; Hass et al., 2004; Taniguchi et al., 2007; Argyros et al., 2008; Heyl et al., 2008; Ishida et al., 2008; Ramireddy et al., 2013). From our results (Figure 5B) it was evident that, the cytokinin biosynthesis genes like *IPT3, IPT5* and *IPT7*, did not show any reduced expression in *rav1*. At specific stages of development, expression of the *IPT* genes was rather higher in *rav1*, compared to wild type. Notably in *rav1*, significantly reduced expression was observed for cytokinin response genes like *ARR1, ARR12* and *ARR6*, compared to wild type. Since expression of the cytokinin response genes did not get upregulated in *rav1* roots, it implied that cytokinin signalling and not biosynthesis was hindered in absence of RAV1. Along with the cytokinin biosynthesis and signalling genes, we also checked expression of *SHY2* and *PIN1* genes that function downstream to cytokinin for regulating auxin signalling in primary root (Tian et al., 2002; Ruzicka et al., 2009). The cell cycle gene *CYCB1;1* known to be expressed in root apical meristems (Doerner et al., 1996; Colón-Carmona et al., 1999; Biancucci et al., 2015) was also tested. Expression of *SHY2*, the Aux/IAA family protein that works as a negative regulator of auxin signalling, had significantly reduced expression in *rav1*, compared to wild type. This suggested that repression of auxin signalling was compromised in *rav1* roots. Correlating to this, expression of the auxin efflux transporter *PIN1*, was found to be increased in *rav1*, compared to wild type. The *CYCB1;1* gene expression was also found to be higher in *rav1*, compared to wild type (Figure 5B). In Arabidopsis, *CYCB1;1* gene encodes the primary mitotic cyclin and ectopic expression of it is known to stimulate cell division in root apical meristems (Doerner et al., 1996; Himanen et al., 2003). Thus, increased expression of *CYCB1;1* gene may have enhanced cell division activities in *rav1* roots. The above results clearly indicate that expression of cytokinin signalling genes is downregulated in absence of RAV1, which consequently affects auxin signalling and transport through altered expression of *SHY2* and *PIN1* genes. This leads to higher expression of *CYCB1;1* gene in *rav1* roots, generating a larger root meristem in the mutant, as clearly evident from our confocal microscopy data (Figure 2). Additionally, we also checked the expression of *CRF1*, an AP2/ERF transcription factor implicated in cytokinin signalling, especially in Arabidopsis roots (Rashotte et al., 2006; Raines et al., 2016). Notably, we found that expression of *CRF1* was significantly higher in *rav1*, compared to wild type (Figure 5B). It is worthy of mention here that high throughput transcriptome analyses from roots of wild type and *rav1* seedlings 14 dag, have revealed significantly higher expression of *CRF1* in the mutant, along with few other genes involved in cytokinin and auxin signalling (Supplemental Figure S4). Interestingly, *CRF1* expression is known to be not affected by cytokinin, and mutation of *CRF1* together with *CRF3, 5* and *6* decrease root meristem size along with reduced expression of several cell cycle related genes (Raines et al., 2016). In consonance, we observed that *rav1* roots which showed higher *CRF1* expression compared to wild type, had larger meristem size and increased expression of at least one root-meristem specific cell cycle gene *CYCB1;1*. To further confirm whether CRF1 has a negative effect on cytokinin signalling during primary root growth, we did transcription analyses with RNA isolated from roots of CRF1-deleted (*crf1*) mutant and CRF1-overexpressing (CRF1OX) seedlings. As shown in Figure 5C, expression of cytokinin response genes like *ARR1* and *SHY2* was upregulated in *crf1* and downregulated in CRF1OX, compared to wild type. On the other hand, *PIN1* whose expression is reduced under cytokinin signalling, was upregulated in CRF1OX and downregulated in *crf1* mutant roots. It is thus imperative to suggest that CRF1 antagonizes cytokinin signalling during primary root development in Arabidopsis. To further correlate, since *CRF1* expression is enhanced in *rav1* roots, RAV1 has a role to play in regulation of *CRF1* expression during primary root growth.

**Figure 5.**
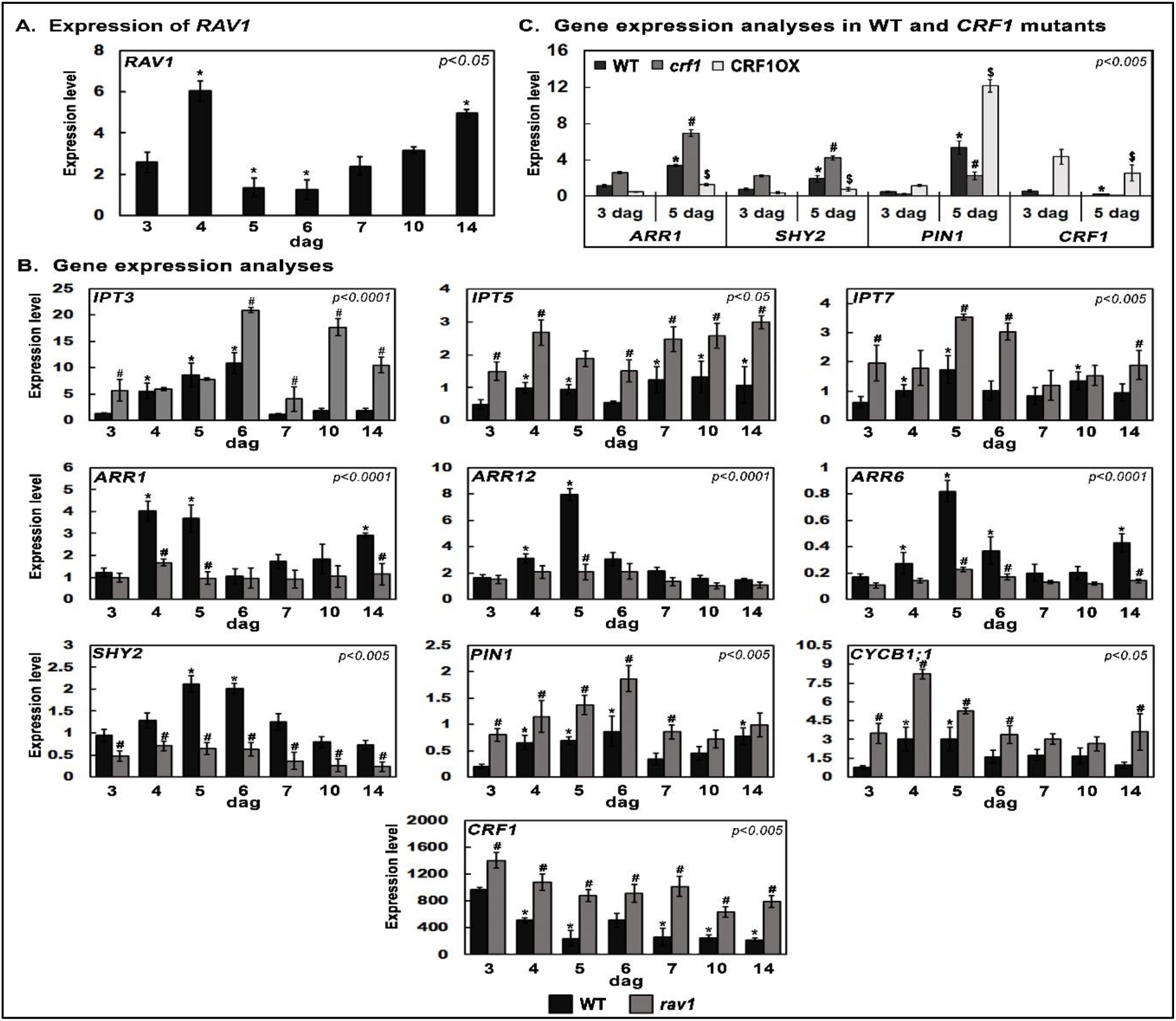
Expression of *RAV1* and comparative expression of cytokinin-responsive genes. Total RNA was isolated from roots of WT, *rav1, crf1* and CRF1OX mutant seedlings at indicated number of days after germination (dag) and subjected to qRT-PCR. **A**. Expression of *RAV1* gene in roots of WT plants. **B**. Expression of cytokinin biosynthesis (*IPT3, IPT5, IPT7*), cytokinin signalling (*ARR1, ARR12, ARR6*), auxin signalling (*SHY2, PIN1*), cell cycle reporter (*CYCB1;1)* and cytokinin response factor (*CRF1*) genes in roots of WT and *rav1. GAPDH* expression was used for normalization. **C**. Comparative gene expression in wild type, *crf1* and CRF1OX roots. All qPCRs were performed in triplicate and error bars represent SE. Anova two-factor with replication method was used to calculate the variance. Variance in between WT and *rav1* is denoted by **#** and variance within different time points of WT is denoted by * on corresponding graphs. Variance between WT and *CRF1* mutants *crf1* and CRF1OX is denoted by **#** and $, respectively, and variance within different time points of WT is denoted by * on corresponding graphs.

### RAV1 negatively regulates expression of *CRF1*

As *CRF1* expression was higher in *rav1* roots, we next wanted to know whether RAV1 can negatively regulate *CRF1* expression. For this, we first attempted a ChIP-qPCR based analyses in wild type plants using anti-RAV1 antibody (Sengupta et al., 2020). As shown in Figure 6A, *in silico* analyses indicated that upstream regulatory region of *CRF1*, had three RAV1-AP2 domain binding sites (CAACA). Hence, primers were designed to PCR amplify one fragment containing the single promoter-proximal CAACA site located around 1.1 kb upstream to ATG (designated as Region I), and another second fragment that contains two more CAACA sites located around 1.8 kb upstream (designated as Region II). As per our findings, prominent RAV1 occupancy was observed in Region I of *CRF1* promoter (Figure 6B). RAV1 occupancy was distinctly higher around 5 and 14 dag, relative to 3 dag. It is worthy of mention here that the RAV1 binding site in Region I has several designated TATA boxes (Berendzen et al., 2006) located in the near vicinity (Figure 6A). The closest one (TATATT) is located only 26 bp away. Thus, binding of RAV1 to the CAACA element in Region I may hinder transcription activation, by occluding the nearby TATA box(es). This further points out to the possibility that RAV1 acts as a repressor of *CRF1* gene expression. No significant recruitment of RAV1 however was observed at promoter distal sites within Region II of *CRF1* (data not shown), or at downstream sites containing no known RAV1 binding element, thus confirming specificity of the ChIP results (Supplemental Figure S5). To further understand whether RAV1 can actually repress *CRF1* promoter activity, a 1.2 kb fragment of *CRF1* corresponding to Region I, containing the single CAACA site was cloned upstream to GUS reporter gene for generating the proCRF1::GUS construct. GUS assay was done in wild type and *rav1* Arabidopsis plants, as described previously (Sengupta et al., 2020). In the *rav1* mutant, where functional RAV1 protein is lacking, there was a significant increase in GUS expression compared to wild type (Figure 6C). This clearly pointed out that absence of RAV1 enhanced *CRF1* promoter activity. To further complement this data, *CRF1* transactivation assay was done in Nicotiana, as well. For this, RAV1 cDNA was cloned under 35S promoter and the resultant construct was co-infiltrated into *Nicotiana benthamiana* leaves, along with proCRF1::GUS construct. Results indicated that GUS activity was significantly reduced when proCRF1::GUS was co-infiltrated with 35S::RAV1 construct, compared to when proCRF1::GUS construct was infiltrated alone (Figure 6D). Presence of RAV1 protein therefore evidently repressed *CRF1* promoter activity. It is thus conclusive from the above results that RAV1 acts as a transcription repressor for *CRF1* gene. To correlate, it can therefore be stated that RAV1 promotes cytokinin signalling in Arabidopsis roots through negative regulation of *CRF1* gene expression.

**Figure 6.**
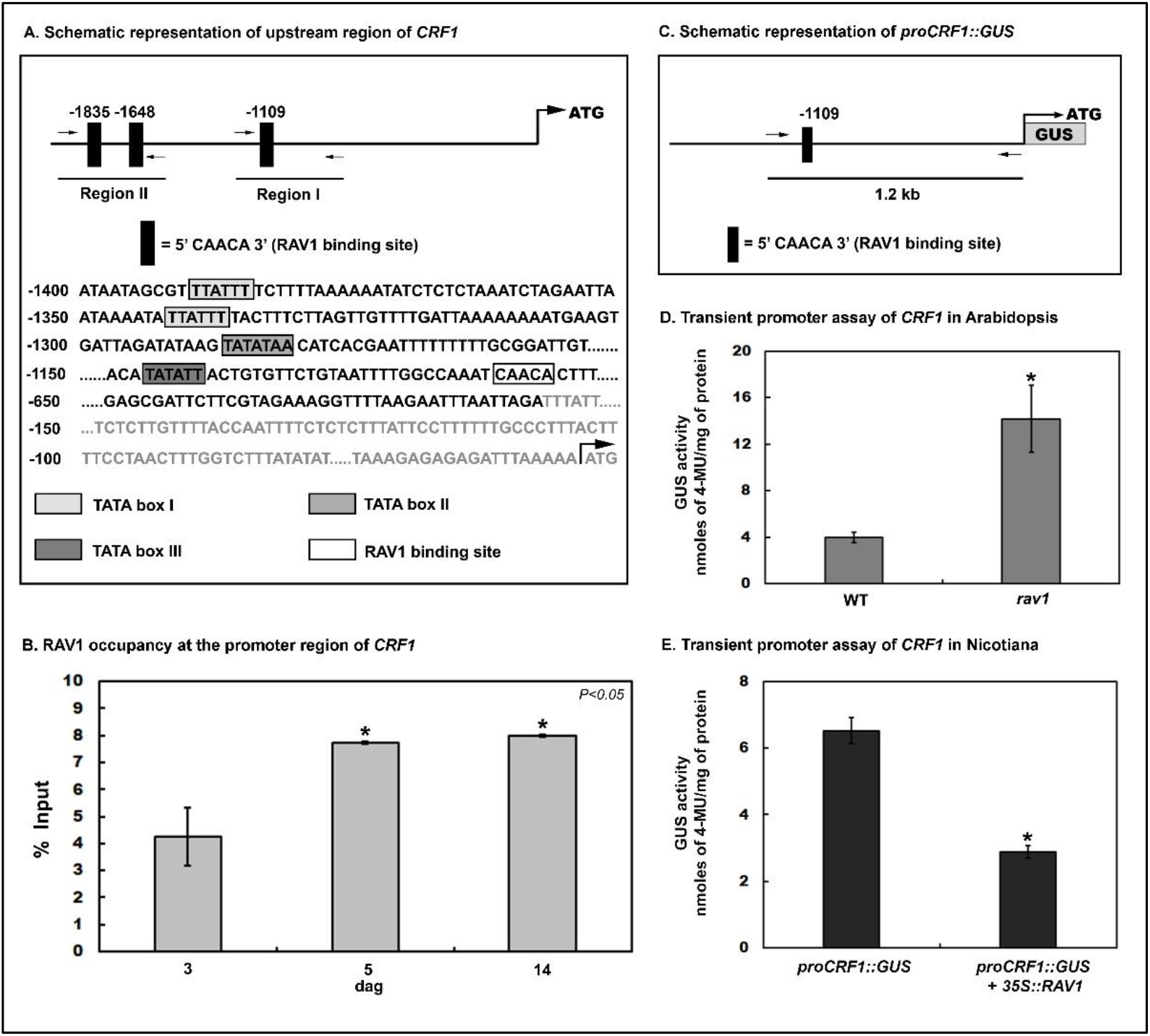
RAV1 negatively regulates *CRF1* promoter activity. **A**. Schematic map of *CRF1* promoter region showing three RAV1 binding sites (CAACA). Bottom panel showing the nucleotide sequence upstream of the transcriptional start site of *CRF1* and location of designated TATA boxes in the vicinity of RAV1 binding site (CAACA) in Region I. **B**. ChIP assay using anti-RAV1 antibody showing RAV1 occupancy at Region I of *CRF1* promoter bearing CAACA sequence, 3, 5, and 14 dag. The data represented is a mean of three independent biological repeats with standard error bars. Statistical significance tested using paired two-tailed *t*-test and results with P value ≤ 0.05 are marked by *. **C**. Schematic representation of *proCRF1::GUS* promoter construct indicating RAV1 binding site (CAACA), used for transient promoter assay of *CRF1*. **D**. Leaves of 6-weeks-old WT and *rav1 Arabidopsis thaliana* plants were infiltrated with either resuspension buffer or *A. tumefaciens* containing *proCRF1::GUS* and were allowed to recover for 48 hrs. The infiltrated leaves were excised and subsequently GUS assay was performed. Values were normalised against resuspension buffer control. Graph represents GUS activity of *CRF1* promoter in WT and *rav1* expressed as nanomoles of 4-MU per mg of total protein. **E**. Leaves of 6-weeks-old *Nicotiana benthamiana* plants were either infiltrated with *proCRF1::GUS* or co-infiltrated with *proCRF1::GUS* and *35S::RAV1* and after 48 hrs recovery infiltrated leaves were excised and used for GUS assay. Graph represents GUS activity expressed as nanomoles of 4-MU per milligram of protein. The represented data in (D) and (E) were mean of three biological replicates over three separate occasions (n ≥ 20 leaves) with standard error bars. Student’s t-test with paired two-tailed distribution was used for statistical analysis and P ≤ 0.005 is denoted by *.

### Cytokinin induces *RAV1* gene expression through ARR1

To further substantiate the role of RAV1 during cytokinin signalling, we next performed gene expression analyses with cytokinin-treated roots of *rav1* and wild type. For this, RNA was isolated from 5-day old *rav1* and wild type seedlings mock-treated or treated with exogenous cytokinin (5 μM 6-BAP) for different time points. As shown in Figure 7A, the data obtained indicate that *CRF1* expression did not alter on cytokinin treatment whether in wild type or *rav1*, although its expression in *rav1* roots remained significantly higher than wild type. Expression of the two type-B genes *ARR1, ARR12* and the type-A gene *ARR6* were significantly upregulated in wild type roots on cytokinin treatment, but failed to show cytokinin induction in *rav1*. Similarly, expression of *SHY2* in wild type showed cytokinin responsive upregulation, but no such cytokinin inducibility could be observed for *SHY2* gene in *rav1*. Interestingly, we observed reduced expression of *PIN1* and *CYCB1;1* in both wild type and *rav1* on cytokinin treatment. Such reduced expression although expected in wild type upon cytokinin treatment, was intriguing for *rav1*. This however did not come as a complete surprise, because with exogenous cytokinin treatment we did observe reversion of *rav1* phenotype, as its primary root length and meristem size became similar to wild type (Figure 2). Thus essentially, upon exogenous cytokinin treatment there must be a pathway that still leads to reduction in expression of *PIN1* and *CYCB1;1*, bypassing the requirement for known cytokinin response genes like the ARRs or SHY2, that consequently can control root meristem size. Future investigation towards this direction may lead to interesting findings. The most conspicuous result of this experiment was cytokinin inducibility of *RAV1*. Significant upregulation of *RAV1* was observed within 30 min of cytokinin treatment in wild type roots (Figure 7A). Collectively, it suggests that RAV1 is transcriptionally upregulated by cytokinin and further promotes cytokinin signalling in primary root through downregulation of *CRF1* and upregulation of several cytokinin-responsive genes like *ARR1, ARR12, ARR6* and *SHY2*.

**Figure 7.**
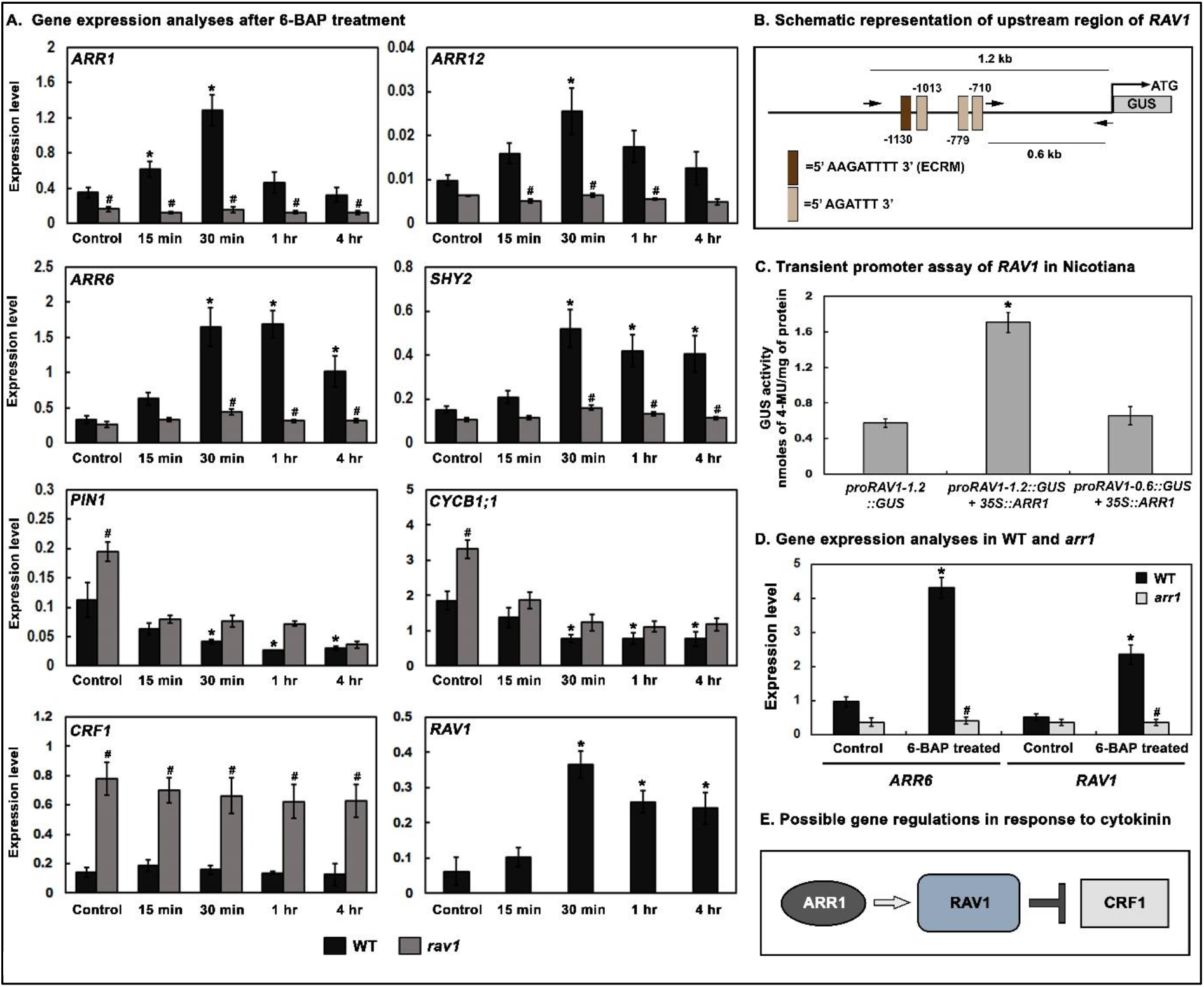
Cytokinin responsive gene expression and upregulation of *RAV1* gene through ARR1. **A**. Total RNA was isolated from roots of WT and *rav1* mutant 5 dag, treated with either 5 μM 6-BAP for 15 min, 30 min, 1 hr, and 4 hr or 0.1 % DMSO control for 4 hr and subjected to qRT-PCR analysis. Anova two-factor with replication method was used to calculate the variance between wild type and *rav1*. Variance in between WT and *rav1* is denoted by **#** and variance within WT samples is denoted by * on corresponding graphs (P ≤ 0.005). **B**. Schematic representation of *RAV1* promoter region showing ARR1 binding sites AGATTT, including an extended cytokinin-response motif (ECRM) AAGATTTT. **C**. Leaves of 6-weeks-old *Nicotiana benthamiana* plants were infiltrated with *proRAV1-1*.*2::GUS* or co-infiltrated with either *proRAV1-1*.*2::GUS* and *35S::ARR1* or *proRAV1-0*.*6::GUS* and *35S::ARR1*. Graph represents GUS activity expressed as nanomoles of 4-MU per milligram of protein. The represented data is a mean of three biological replicates over three separate occasions (n ≥ 20 leaves) with standard error bars. Student’s t-test with paired two-tailed distribution was used for statistical analysis and P ≤ 0.005 is denoted by *. **D**. Gene expression analyses in WT and *arr1*, treated with either 5 μM 6-BAP or 0.1 % DMSO control for 30 min. Anova two-factor with replication method was used to calculate the variance between wild type and *arr1*. Variance in between WT and *arr1* is denoted by # and variance within WT samples is denoted by * on corresponding graphs (P ≤ 0.05). **E**. A schematic of possible gene regulation as a part of cytokinin signalling in Arabidopsis, depicting a pathway where ARR1 positively regulates *RAV1* gene expression while RAV1 negatively regulates *CRF1* gene expression.

As *RAV1* expression is induced in response to cytokinin, it therefore warranted the need to understand how the hormone upregulates *RAV1* gene expression. As shown in Figure 7B, *in silico* analyses revealed that RAV1 upstream region has several conserved ARR1 binding sites AGAT(T/C)(T/C) in the promoter region, including an extended cytokinin-response motif (ECRM) AAGAT(T/C)TT (Sakai et al., 2001; Ramireddy et al., 2013; Xie et al., 2018). Transient promoter assay was hereafter done to check whether ARR1 can transactivate *RAV1* expression. For this, as shown in Figure 7B, a 1.2 kb region in the *RAV1* promoter containing the ARR1 recognition sites was cloned upstream to GUS reporter gene, designated hereafter as proRAV1-1.2::GUS. The DNA binding domain along with the transactivation domain (Sakai et al., 2000) of ARR1 was cloned under a 35S promoter (35S::ARR1). GUS assay was done following infiltration of either proRAV1-1.2::GUS promoter construct alone or along with 35S::ARR1 into *Nicotiana benthamiana* leaves. Results indicated that GUS activity was significantly higher in presence of ARR1, compared to when proRAV1-1.2::GUS construct was infiltrated alone (Figure 7C). To further confirm the specificity of ARR1 mediated transactivation of *RAV1*, GUS assay was done with a proRAV1-0.6::GUS construct, bearing a 0.6 kb promoter region of *RAV1* without any ARR1 binding site. Co-infiltration of proRAV1-0.6::GUS construct was done into *Nicotiana benthamiana* leaves along with 35S::ARR1. As evident from the results, GUS activity of proRAV1-0.6::GUS construct in presence of ARR1 was almost similar to activity of proRAV1-1::GUS construct alone (Figure 7C).

To further confirm that ARR1 mediates *RAV1* gene expression in response to cytokinin, we checked *RAV1* expression in *arr1* mutant (NASC ID - N6971) (Argyros et al., 2008), with or without cytokinin treatment. As evident from our results, no significant upregulation of *RAV1* was observed in *arr1* roots in response to cytokinin, compared to wild type, indicating that cytokinin-induced *RAV1* expression requires functional ARR1 (Figure 7D). To further substantiate this result, we also checked expression of *ARR6* gene, the direct target of ARR1 in cytokinin signalling. As shown in our results, unlike wild type, no cytokinin-induced increase in *ARR6* expression was observed in *arr1* roots. Collectively, these results indicate that similar to *ARR6, RAV1* gene expression is also positively regulated by ARR1, as a part of cytokinin signalling.

To sum up, as shown in Figure 7E, gene expression and promoter assay results obtained thus far suggests that ARR1 transactivates *RAV1* gene expression, and RAV1 negatively regulates *CRF1* gene expression, in response to cytokinin signalling.

### *RAV1* is a secondary response gene in cytokinin signalling

ARR1, typically a type-B ARR, is considered a primary cytokinin responsive gene that is involved in expression of further downstream genes during cytokinin signalling (Sakai et al., 2000; Sakai et al., 2001; Mason et al., 2005). We next wanted to check that whether ARR1-mediated expression of *RAV1* is a primary response similar to *ARR6* expression, or part of a secondary response, in context to cytokinin signalling. To check this, we did transcription analyses with roots of wild type seedlings that have been pre-treated with translation inhibitor cycloheximide (CHX), followed by treatment without or with cytokinin for different time periods. As shown in Figure 8, it was observed that cytokinin treatment could upregulate expression of known primary cytokinin responsive genes like *ARR1, ARR12* and *ARR6*, even after cycloheximide treatment. Expression of *SHY2*, which is known to get induced downstream to ARR factors (Moubayidin et al., 2009; Moubayidin et al., 2010), did not get upregulated by cytokinin upon cycloheximide treatment, confirming that it functions as a secondary response gene. Expression of the other secondary response genes *PIN1* and *CYCB1;1* also did not change upon cytokinin treatment in presence of cycloheximide. Additionally, expression of *CRF1* which is not induced cytokinin showed no difference on treatment with cycloheximide. Most notably, expression of RAV1, which was found to be significantly upregulated by cytokinin, showed no cytokinin-mediated upregulation in presence of cycloheximide. This strongly suggested that *RAV1* acts as a secondary cytokinin response gene that is upregulated by ARR1 in presence of cytokinin. RAV1 further works towards transcription regulation of several downstream genes including *CRF1*, to promote cytokinin signalling in primary root.

**Figure 8.**
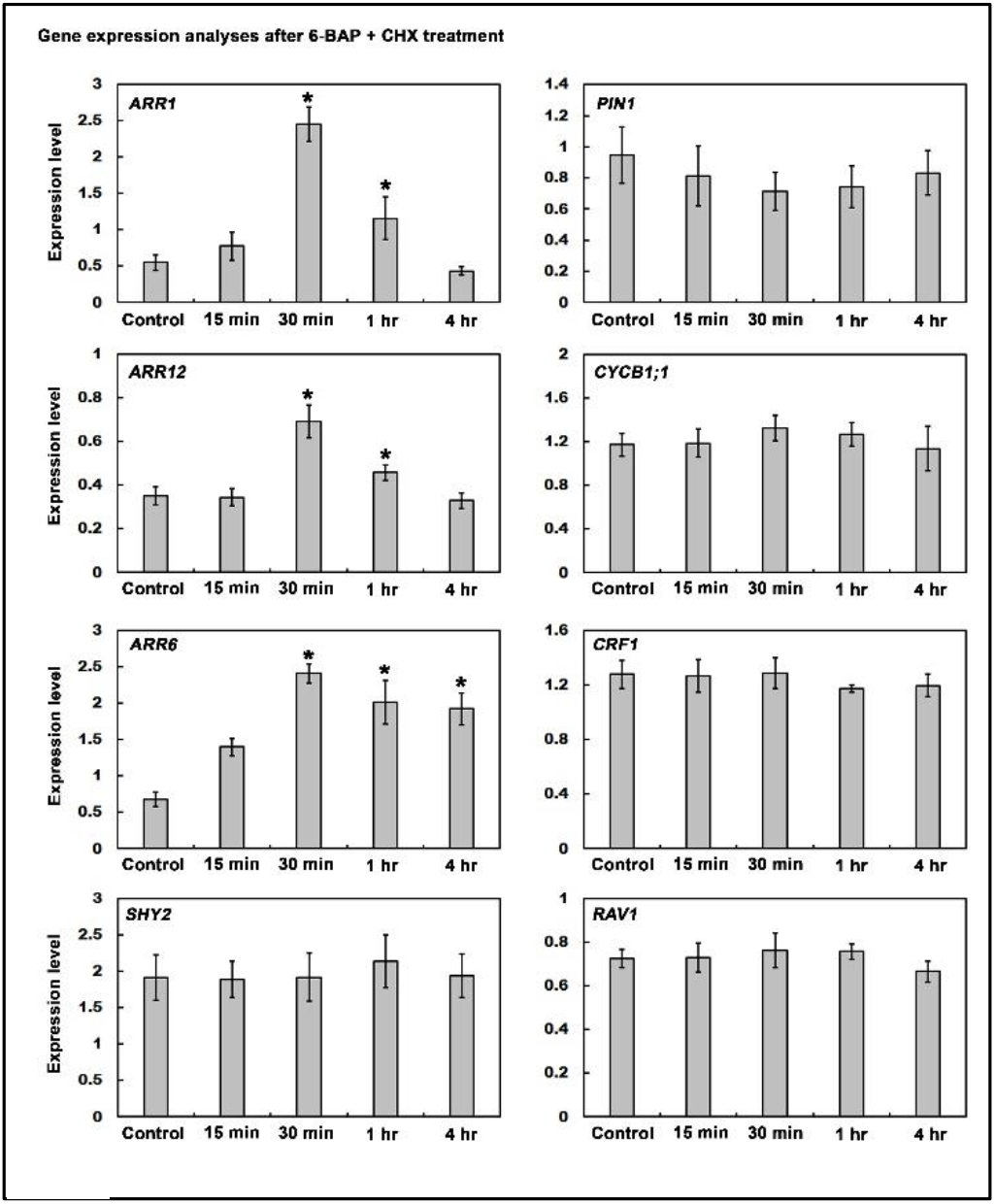
Primary and secondary gene expression analyses in response to cytokinin. WT seedlings 5 days after germination, were pre-treated with 50 μM Cycloheximide (CHX) for 30 min and then 5 μM 6-BAP + 50 μM Cycloheximide (CHX) for 15 min, 30 min, 1 hr, and 4 hr as described in Materials and methods. Total RNA was extracted from the roots at indicated time point and subjected to qRT-PCR. The data represented are mean of three independent biological repeats with standard error bars. Statistical significance tested using paired two-tailed student’s t-test and results with P ≤ 0.005 are marked by *.

## Discussion

Primary root growth at the post-embryonic stage is a dynamic process and involves complex interplay of several hormones, acting in concert or in opposition. Previous reports have indicated that *rav1* mutant seedlings have increased primary root growth compared to wild type, under control conditions, as well as under dehydration stress (Sengupta et al., 2020). The mutant phenotype was intriguing and, in our efforts to get a mechanistic insight we observed that despite delayed germination, the primary root of *rav1* mutant overgrew that of wild type by 4 days after germination. The root length differences increased and reached to a level where *rav1* roots were longer by several centimetres compared to wild type, within two weeks after germination. Subsequent microscopic observations conclusively indicated that the *rav1* mutant roots had a larger meristem with increased cell numbers, compared to wild type. Arabidopsis primary root is prominently classified into four zones (Verbelen et al., 2006), and each zone is characteristically an outcome of a complex web of hormonal crosstalk. The dynamism observed in root growth is distinctly dependent on the balance struck between division and differentiation of the root meristem cells, an event prominently helmed by the interplay of two hormones cytokinin and auxin. It is widely known that cytokinin acts at the transition zone to restrict auxin distribution and effectively regulate root meristem size (Dello Ioio et al., 2007; Dello Ioio et al., 2008; Dello Ioio et al., 2008; Ruzicka et al., 2009; Moubayidin et al., 2010; Šimášková et al., 2015). The fact that similar to the cytokinin response mutant *arr1* (Sakai et al., 2001) the phenotype of *rav1* could be reversed with exogenous cytokinin treatment, indicated that impaired cytokinin action was the underlying cause for the observed *rav1* root morphology. Findings from the microfluidics-based experiment further confirmed this understanding. It was evident that cytokinin when applied specifically to *rav1* root tip within 4 dag could reinstate root meristem to a size that is comparable to wild type grown under control conditions. Use of microfluidics platform to deliver hormonal stimuli to Arabidopsis root, was first reported by Ismagilov and his group (Meier et al., 2010). Similar studies have since then been done to check the effect of heterogeneous environment on root growth, gene expression and root-microbe interactions (Busch et al., 2012; Grossmann et al., 2012; Massalha et al., 2017; Park et al., 2017; Stanley et al., 2018). However, no research endeavour thus far, has compared mutant and wild type root growth in the microfluidics platform as reported in this work. Moreover, treating predefined region of the *rav1* mutant root with cytokinin, at a specific spatial resolution, and resultant alteration of the root meristem size is an unprecedented approach. The inference drawn from the above results is crucial and clearly indicate that absence of RAV1 impairs cytokinin signalling in primary root of Arabidopsis and the longer root phenotype of *rav1* can be reverted with cytokinin treatment. Similar effect was observed by germinating *rav1* in presence of auxin transport inhibitor NPA. As auxin transport and distribution was significantly enhanced in *rav1* primary roots, the effect of NPA-mediated inhibition and reduction in root growth was much more prominent in the mutant, compared to wild type. It thus further confirmed that absence of RAV1, altered cytokinin signalling and effectively auxin transport and distribution during primary root growth in Arabidopsis. Till date, RAV1, an AP2/ERF family transcription factor although implicated in ABA, ethylene and brassinosteroid signalling (Hu et al., 2004; Ikeda and Ohme-Takagi, 2009; Feng et al., 2014; Sengupta et al., 2020), has never been directly related to cytokinin signalling and its downstream effects. A previous work with suspension culture has shown that, *RAV1* expression in *det2* mutant that has defect in brassinosteroid biosynthesis, shows marginal downregulation in presence of zeatin. The authors however, have not established any correlation between RAV1 and cytokinin signalling in wild type cells (Hu et al., 2004). Our present findings show for the first time that RAV1 plays a distinct role in cytokinin signalling pathway during primary root development.

Further efforts to unravel the molecular mechanism of RAV1 mediated cytokinin signalling revealed that absence of RAV1 did not impede transcription of cytokinin biosynthesis genes in Arabidopsis primary root. RAV1 downstream regulon rather comprised of the cytokinin signalling genes like *ARR1, ARR12* and *ARR6*, as expression of these genes were severely downregulated in the *rav1* mutant. Interestingly, in *rav1* roots exogenous cytokinin treatment could not upregulate these cytokinin response genes, unlike wild type. It thus became evident that impaired cytokinin signalling was the primary cause of larger root meristem in the *rav1* mutant. It has been previously reported that *arr1arr12* double mutant seedlings show longer primary root growth compared to wild type (Dello Ioio et al., 2007), which is similar to our observation on *rav1* mutant phenotype. A logical correlation could be that, longer primary root growth observed in *rav1* mutant relative to wild type, is a manifestation of reduced expression of *ARR1* and *ARR12* genes in the mutant. Additionally, expression of *SHY2*, the auxin signalling repressor was also significantly downregulated in *rav1* mutant, compared to wild type. SHY2 an Aux/IAA family member is known to get activated by cytokinin in root and impairs expression of downstream *PIN1*, the auxin transporter molecule (Tiwari et al., 2001; Dello Ioio et al., 2007; Taniguchi et al., 2007; Dello Ioio et al., 2008; Dello Ioio et al., 2008). In concurrence, we found that lower levels of *SHY2* expression in *rav1* roots is accompanied by higher levels of *PIN1* expression and broader auxin distribution, compared to wild type. The *SHY2* and *PIN1* gene expression data complements the observation that the longer root phenotype and larger meristem size in the mutant was reversed, when grown in presence of auxin transport inhibitor, NPA. Therefore, to sum up, in absence of RAV1 auxin transport and distribution is altered in Arabidopsis primary root. Such differential auxin maxima in the *rav1* primary root enhances its root meristem size and root length, compared to wild type. As cytokinin regulates auxin transport for maintaining auxin maxima towards the root tip and for determining root meristem size, impaired cytokinin signalling in *rav1* root, effectively fails to regulate auxin distribution and in turn cannot control root meristem size in the mutant.

The CRFs, belonging to the AP2/ERF family of transcription factors, have recently been implicated to play diverse role in cytokinin signalling, including in roots (Jeon et al., 2016; Raines et al., 2016). *In silico* analyses revealed that amongst the CRFs known to function in roots, *CRF1* and *CRF3* have RAV1 binding sites (CAACA), in their respective upstream regulatory region. *CRF3* has been primarily implicated in regulating lateral root development (Jeon et al., 2016). In trying to relate *CRF1* with primary root development we found that its expression is higher in *rav1* roots, compared to wild type. This observation was important and indicated that post-embryonic root development in Arabidopsis requires reduction in *CRF1* expression. Evidently, RAV1 functions as a transcription repressor of *CRF1* gene through recruitment to the *CRF1* promoter region. Reducing expression of *CRF1*, is thus the possible pathway through which RAV1 promotes cytokinin signalling. In Arabidopsis, cytokinin perception and cytokinin-mediated differentiation at the transition zone reaches maxima around 4-5 days after germination, until then the root meristem grows. During this time high levels of *ARR1* expression occurs and sets a balance between cell division and differentiation which finally regulates the root meristem size (Dello Ioio et al., 2007; Dello Ioio et al., 2008; Dello Ioio et al., 2008). Interestingly, this seems to be the time window when cytokinin signalling peaks for regulating root meristem size, through downregulation of *CRF1* and upregulation of *ARR1*. The role of CRFs in cytokinin signalling is complex, but it has been shown that they tend to antagonize the signalling by expression regulation of several genes including Type-A ARRs, that are negative regulators of cytokinin response (Rashotte et al., 2006; Raines et al., 2016). Our gene expression analyses with roots from *crf1* and *CRF1ox* seedlings further indicated that in absence of *CRF1*, expression of cytokinin-response genes like *ARR1* and *SHY2* is significantly increased, while that of *PIN1* is reduced, compared to wild type. This indicates a role of CRF1 in negatively regulating cytokinin signalling. Taken together, it is pertinent to summarize that RAV1 represses *CRF1* expression to promote cytokinin signalling during primary root development. Increased expression of *CRF1* in *rav1* roots negatively impacts cytokinin signalling and consequent control over root meristem size. The fact that cytokinin responsive genes like *ARR1, ARR1*2, *ARR6* and *SHY2* could not get upregulated in *rav1* roots by exogenous cytokinin treatment, further emphasized that it was impaired cytokinin response and not impaired cytokinin biosynthesis that affected *rav1* primary roots.

The most intriguing finding from our study was cytokinin inducibility of *RAV1* gene expression and the fact that the *RAV1* promoter has several conserved and extended (ECRM) ARR1 binding sites. The fact that ARR1 could transactivate *RAV1* promoter was quite evident from the transient promoter assay results indicating that *RAV1* functions downstream to ARR1 during cytokinin signalling in Arabidopsis primary root. This was further confirmed when cytokinin induced upregulation of *RAV1* expression was not observed in the roots of *arr1* mutant. However, unlike *ARR6* which is a primary response gene and a direct target of ARR1 during cytokinin signalling, *RAV1* is a secondary cytokinin-responsive gene. Similar to *SHY2*, which is a known secondary response gene in cytokinin signalling, expression of *RAV1* could not be induced directly by cytokinin in presence of protein translation inhibitor cycloheximide. Thus, *RAV1* expression regulation mediated by ARR1 is a branch of cytokinin signalling cascade that works towards repression of *CRF1* and in turn augments the hormone signalling in Arabidopsis root.

To conclude, as represented in Figure 9, ARR1-mediated *RAV1* expression regulation is directed towards repression of *CRF1* gene. Such a string of expression regulation of transcription factors is critical for efficient transduction of cytokinin signalling in primary roots. These upstream events further enables SHY2 mediated repression of auxin transporter *PIN1* and consequently regulates auxin distribution, such that root meristem size is controlled. Thus, length control of Arabidopsis primary root, is an effect of intricate crosstalk between auxin and cytokinin signalling, that is critically regulated by RAV1. From such findings on role of RAV1 in regulating primary root growth, it becomes apparent as to why the ABA responsive transcription factor ABI3, reduces *RAV1* gene expression as a part of dehydration stress response (Sengupta et al., 2020). The reprogramming of primary root length regulation during water stress management is a critical parameter that determines the stress response efficiency of any plant system. Regulation of *RAV1* itself and the downstream events that are mediated by RAV1 must be one of the important criteria in this aspect.

**Figure 9.**
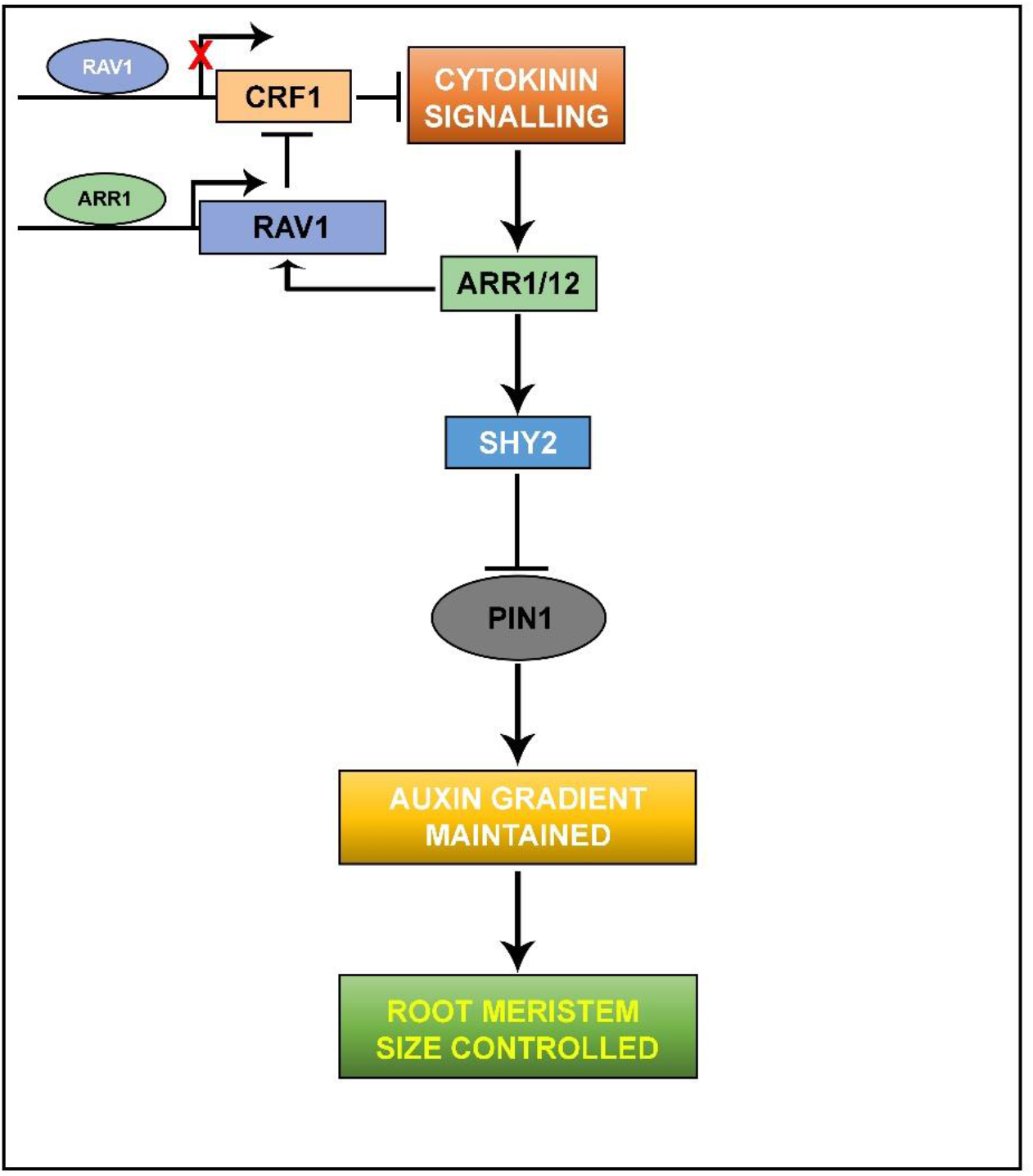
A schematic of possible pathway illustrating involvement of RAV1 as a part of cytokinin signalling that regulates root meristem development in Arabidopsis. In response to cytokinin, ARR1 transactivates *RAV1* and RAV1 negatively regulates *CRF1* to promote cytokinin signalling in Arabidopsis root. Cytokinin signalling further promotes upregulation of *SHY2* expression downstream to ARR1, eventually leading to downregulation of *PIN1* expression. Such regulation of gene expression effectively results in maintenance of auxin gradient in Arabidopsis primary root, and controls root meristem size.

## Materials and Methods

### Plant materials and growth conditions

All genotypes are in the *Arabidopsis thaliana* ecotype Columbia (*Col-0*) background. The following Arabidopsis seed stocks were used: *rav1* (referred herein as “*rav1*”, NASC stock ID-N420832, GABI-KAT, Bielefeld University, Bielefeld, Germany) (Sengupta et al., 2020), another *rav1* mutant (referred herein as “*rav1-1*”, NASC stock ID-N655012, NASC, The University of Nottingham, Loughborough, UK), *arr1* (NASC stock ID-N6971, NASC) (Argyros et al., 2008). *CRF1* mutant lines (*crf1*, GABI_068G09 and CRF1OX) were generously provided by Dr. Joseph J. Kieber at Department of Biology, University of North Carolina, Chapel Hill, NC 27599, USA. The mutant lines were confirmed by PCR using primers as listed in Supplemental Table S1. The seeds were surface sterilized (Bedi et al., 2016; Bedi and Nag Chaudhuri, 2018; Sengupta et al., 2020) and vernalized at 4 °C for 2 days in darkness and then plated on Murashige and Skoog (MS) medium. For morphological analyses, seedlings were grown vertically (22 °C, 16 h light/8 h dark) for the indicated number of days. *Nicotiana benthamiana* seeds were directly sown on potting soil (Soilrite Mix, Keltech Energies Ltd., India) and grown at 25 °C in the plant growth room.

### Promoter and gene cloning

For cloning of DNA binding (DBD) along with transactivation domain (TAD) of *ARR1* (1374 bp) and full-length coding sequence (CDS) of *RAV1* (1034 bp) and for promoter::GUS reporter fusion construct, (*proRAV1-0*.*6 kb, proRAV1-1*.*2 kb, proCRF1-1*.*2 kb* and *proPIN1-1*.*3 kb*) sequences of the genes were amplified from cDNA and genomic DNA of Arabidopsis *Col-0* plants respectively. DR5::GFP plasmid construct was generously gifted by Prof. Takuya Suzaki, University of Tsukuba, Japan and Dr. Praveen Verma, National Institute of Plant Genome Research. Details of the primers used for cloning and plasmid constructions are listed in Table S1. The amplified and purified PCR product was first cloned into a gateway pENTR/d-TOPO entry vector (Invitrogen, Life Technologies) and were subsequently cloned in pKGWFS7. For over-expression under *CaMV 35S* promoter (35S::ARR1 and 35S::RAV1), *ARR1* (DBD+TAD) and *RAV1* (CDS) entry clones were recombined with pK7WG2D. Promoter fragment of *proCRF1-1*.*2 kb* and *proPIN1-1*.*3 kb* was amplified and ligated into TA vector (pGEM®-T Easy Vector, Promega). The proCRF1::GUS and proPIN1::GFP was constructed by replacing *CaMV 35S* promoter of pCAMBIA1304 with *proCRF1-1*.*2kb* and *proPIN1-1*.*3kb*, respectively. All the recombinant plasmids and pCAMBIA1304 were subsequently transferred in either *Agrobacterium tumefaciens* strain LBA4404 *or Agrobacterium rhizogenes* strain R1000.

### Hormone and inhibitor treatment

Arabidopsis seeds were germinated vertically on MS media plates containing either 6-BAP (6-Benzylaminopurine, HiMedia Laboratories Pvt Ltd, India) (conc. 1 nM, 10 nM 50 nM or 100 nM) or 0.1 % DMSO (control) and then photographed 5 days after germination (dag). For *N*-1-naphthylphthalamic acid (NPA) (Sigma Aldrich, USA) treatment, Arabidopsis seeds were germinated on MS medium supplemented with either 0.1 % DMSO or 50, 100, 500 nM NPA, respectively.

### Microfluidics

For analysing the real-time root growth of *Arabidopsis thaliana*, a Plant Root Microfluidic System (PRMS) was developed (Figure 3, Supplemental Figure S2). The system consists of four fabricated microfluidic plant root channels having a width and height of 800 μm and a length of 5 cm. Individual channels were connected in parallel to assemble the final device (PRMS) as shown in Figure S2. The channels are fabricated with polydimethylsiloxane (PDMS, Sylgard-184, Sigma Aldrich, USA) substrate using the wire-drawing method (Jia et al., 2008; Song et al., 2010). All channels in PRMS were connected to a common header through individual inlet ports, while the outlet ports were extended to the collection sump using connectors. The entire setup was placed in a closed chamber to avoid contamination during plant growth and was placed in a plant growth room (22 °C, 16 h light /8 h dark). Syringe pumps (Cole-Parmer, USA) containing 20 ml syringe (NIPRO, India) were used to supply 1/2-strength liquid MS media in the channels through the common header. The media flow rate was set as 0.8 ml/hr, and each channel pertaining to the current flow configuration was designed to get a flow rate of 0.2 ml/hr. The selected flow rate of the media was sufficient to displace the fluid continuously through the channel without creating severe hydrodynamic stress on the root surface. Surface sterilized Arabidopsis seeds were germinated in 5 mm long pipette tip cone embedded in solid MS medium (0.8 % agar and 3 % sucrose, pH 5.8) (Grossmann et al., 2012). 3 dag the pipette tip cone containing seedlings were inserted into the perpendicular slot positioned 1.5 cm away from the inlet port of the PRMS such that growing roots can conveniently enter into the channel. PRMS channel was supplied with 1/2-strength MS media for 12 hours through the inlet port. Subsequently, for hormone treatment WT and *rav1* plants were supplied with 1/2-strength MS media supplemented with 6-BAP (5 μM) or 0.1 % DMSO (control) for indicated time points. Each experiment was performed on three biological replicates over three separate occasions.

### Measurement of root length and meristem cell number

For morphological analysis, root lengths were measured using ImageJ software. The number of root meristem cells were measured as described (Casamitjana-MartÍnez et al., 2003; Dello Ioio et al., 2007) using ImageJ software. Error bars were obtained based on the average of n ≥ 30 seedlings in three biological replications. Student’s t-test with paired two-tailed distribution was used for statistical analysis and P ≤ 0.05 was denoted by either an * or # on corresponding graphs.

### Hairy root transformation

Hairy root transformation of Arabidopsis was done as described previously (Limpens et al., 2004; Bandaranayake and Yoder, 2018). Briefly, 2-day and 7-day old Arabidopsis seedlings were used for co-cultivation with *Agrobacterium rhizogenes* R1000 harbouring either proPIN1::GFP or DR5::GFP or 35S::GFP (pCAMBIA1304). Seedlings were excised at the root-shoot junction and the hypocotyl was coated with *A. rhizogenes* culture. The inoculated seedlings were placed on sterile filter paper in a petri plate containing MS medium supplemented with 3 % sucrose and 400 μM acetosyringone (HiMedia Laboratories Pvt Ltd, India). 10 days after co-incubation plants were observed under fluorescence microscope.

### Microscopic analyses

For confocal microscopy, roots were dipped in propidium iodide (PI) solution (10 µg/ml) for 30 secs to stain the root cell wall (Leica SP8 confocal microscope, Ex-535 nm and Em-610 to 640 nm). Imaging of hairy root transformed plants was done using the Leica fluorescence microscope DM4B equipped with Leica DFC3000 camera with GFP filter settings. For capturing real-time images of the growing roots in PRMS, the channels were placed under the Leica DMI 3000M inverted microscope equipped with a high-speed camera (Phantom, USA and Model: Miro-LAB 320). Images of the roots were taken at intervals of 1 hr, 2 hr, 4 hr and 5 hr to observe growth dynamics and changes in meristematic zone.

### RNA isolation for gene expression analysis

For gene expression analysis under control conditions, roots were harvested at indicated number of days. Cytokinin treatment was carried out by treating 5 dag seedlings (WT, *rav1*, and *arr1*) with either 5 μM 6-BAP for indicated time points or 0.1 % DMSO control for 4 hr in constant light with mild shaking at 22 °C. For cycloheximide (CHX) treatment (D’Agostino et al., 2000; Rashotte et al., 2003) 5 dag seedlings were first immersed in liquid MS media with 50 μM CHX (HiMedia Laboratories Pvt Ltd, India) for 30 min with gentle shaking. 6-BAP was added to a final concentration of 5 μM and plants were incubated for 15 min, 30 min, 1 hr, and 4 hr. CHX pre-treated seedlings were immersed in liquid MS with DMSO 0.1 % for 4 hr at 22 °C for mock treatment. Total RNA was extracted using the RNASure® Plant Mini Kit (Genetix brand, NP-84905) as described by the manufacturer, and cDNA was prepared from the total RNA with RevertAid Reverse Transcriptase (Thermo Scientific™, EP0441). The synthesized cDNA was used for gene expression analyses by qPCR using gene-specific primers (Supplemental Table S1).

### Fluorometric *GUS* assay

Fluorometric *GUS* assay was performed as described previously (Bedi and Nag Chaudhuri, 2018; Sengupta et al., 2020). Values were normalized against resuspension buffer control. Total protein concentration was measured by the Bradford method at 595 nm in spectrophotometer (Hitachi U-2900 UV/VIS spectrophotometer) with Bovine serum albumin (BSA) as standard. GUS activity was expressed as nanomoles of 4-MU produced per mg of protein. GUS measurements were obtained from at least twenty leaves (n ≥ 20) on three biological replicates over three separate occasions. Error bar represents SE and significant difference were calculated using a paired Student’s t-test with two-tailed distribution. P ≤ 0.05 was denoted by * on corresponding graphs.

### Chromatin Immunoprecipitation (ChIP) assay

Nuclei isolation and subsequent ChIP assay was done as previously described (Bedi et al., 2016; Sengupta et al., 2020). The purified DNA pellets were suspended in TE and used for qPCR with specific primer sets (Supplemental Table S1). As shown in Supplemental Table S1, ChIP-PCR for negative control was done with primers specific to regions in *CRF1, EIN2* and *ACT1* loci that contain no known RAV1 binding sites and analysed in 2.5 % agarose gel.

### Realtime PCR and data analysis

For qRT-PCR based gene expression analyses, diluted cDNA was used and *GAPDH* was used as the internal control. The mRNA levels for each target genes were calculated using the mean Ct values which were used to determine ΔCt (Ct[sample]-Ct[internal control]) and finally, the expression level was determined using 2^(-ΔCt)^. For ChIP experiment, the C_T_ values of the input samples were extrapolated to 100% as Input samples were 10% of the cell extract, and the C_T_ values of IP (immunoprecipitated DNA) were plotted as a percentage of Input using the formula 100*2^(Ct^Input^-Ct^IP). All PCRs were performed in triplicate and paired Student’s t-test with two-tailed distribution was used for statistical analysis. P ≤ 0.05 was denoted by an asterisk (*) on corresponding graphs. Anova two-factor with replication method was used to calculate the variance between wild type and *rav1*. P value was denoted by a hash (#) on corresponding graphs.

## Acknowledgement

We sincerely acknowledge, Department of Mechanical Engineering, Indian Institute of Technology, Guwahati, India for providing the infrastructure for conducting all microfluidics related experiments mentioned in the present work. We are grateful to Prof. Joseph J. Keiber, University of North Carolina, Chapel Hill, USA for sharing the seeds of CRF1 deletion mutant and CRF1 overexpression lines. Our sincere thanks to Prof. Takuya Suzaki, School of Life and Environmental Sciences, University of Tsukuba, Japan for his kind gift of the DR5::GFP plasmid construct. We acknowledge the Confocal Microscopy facility of Bose Institute, Kolkata for allowing us to perform necessary experiments. Our sincere thanks to Prof. Maitrayee DasGupta for critical reading of the manuscript.

We sincerely acknowledge the funding support by Science and Engineering Research Board (SERB), India (Scheme #CRG/2018/000461) to Dr. Ronita Nag Chaudhuri, Department of Biotechnology, St. Xavier’s College, Kolkata; CSIR fellowship [08/548(0010)/2021-EMR-I] to Saptarshi Datta; by Department of Science and Technology, India [DST-FIST Grant No. SR/FST/COLLEGE-014/2010(C)] and by Department of Biotechnology, India Grant No. BT/INF/22/SP41296/2020 to St. Xavier’s College, Kolkata.

## Authors Contribution

RNC, DM and SD designed the experiments; DM and SD performed the experiments; RNC, DM and SD analysed the results. Microfluidics based experiments designed by PKM and RNC; the experiments were performed by GR, SD and DM. Manuscript written by RNC, DM and SD; writing of sections pertaining to Microfluidics related experiments contributed by PKM and GR. All the authors approve of the final version of the manuscript.

