## Supplementary data for "RAV1 mediates cytokinin signalling for regulating primary root growth in Arabidopsis"

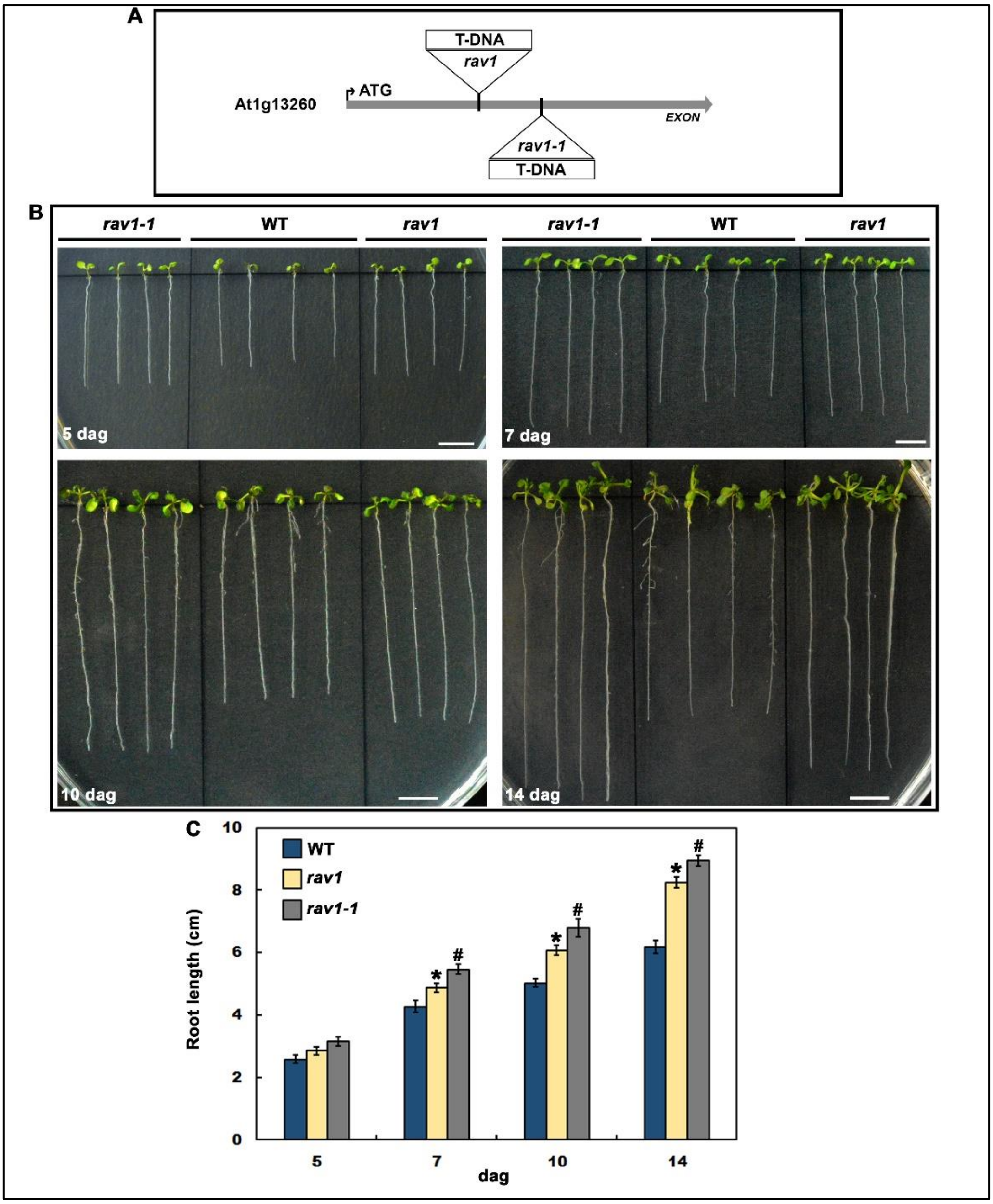


**Figure S1. Both the T-DNA insertion lines of *RAV1* exhibit longer root length phenotype. A.** Schematic showing T-DNA insertions in the respective *rav1* and *rav1-1* mutants. **B.** Morphology of primary root growth of wild-type (WT), *rav1* and *rav1-1* mutants grown vertically on MS medium for 5, 7, 10 and 14 days after germination (dag). Scale bars = 1 cm. **C.** Primary root length measurements of WT, *rav1*and *rav1-1* mutants over time. The data represent the mean of three replicate experiments (n ≥ 20) with standard error bars. Student's t-test with paired two-tailed distribution was used for statistical analysis and P ≤ 0.005 was considered to be significant. Variance in between WT and *rav1* is denoted by * and variance in between WT and *rav1-1* is denoted by #.

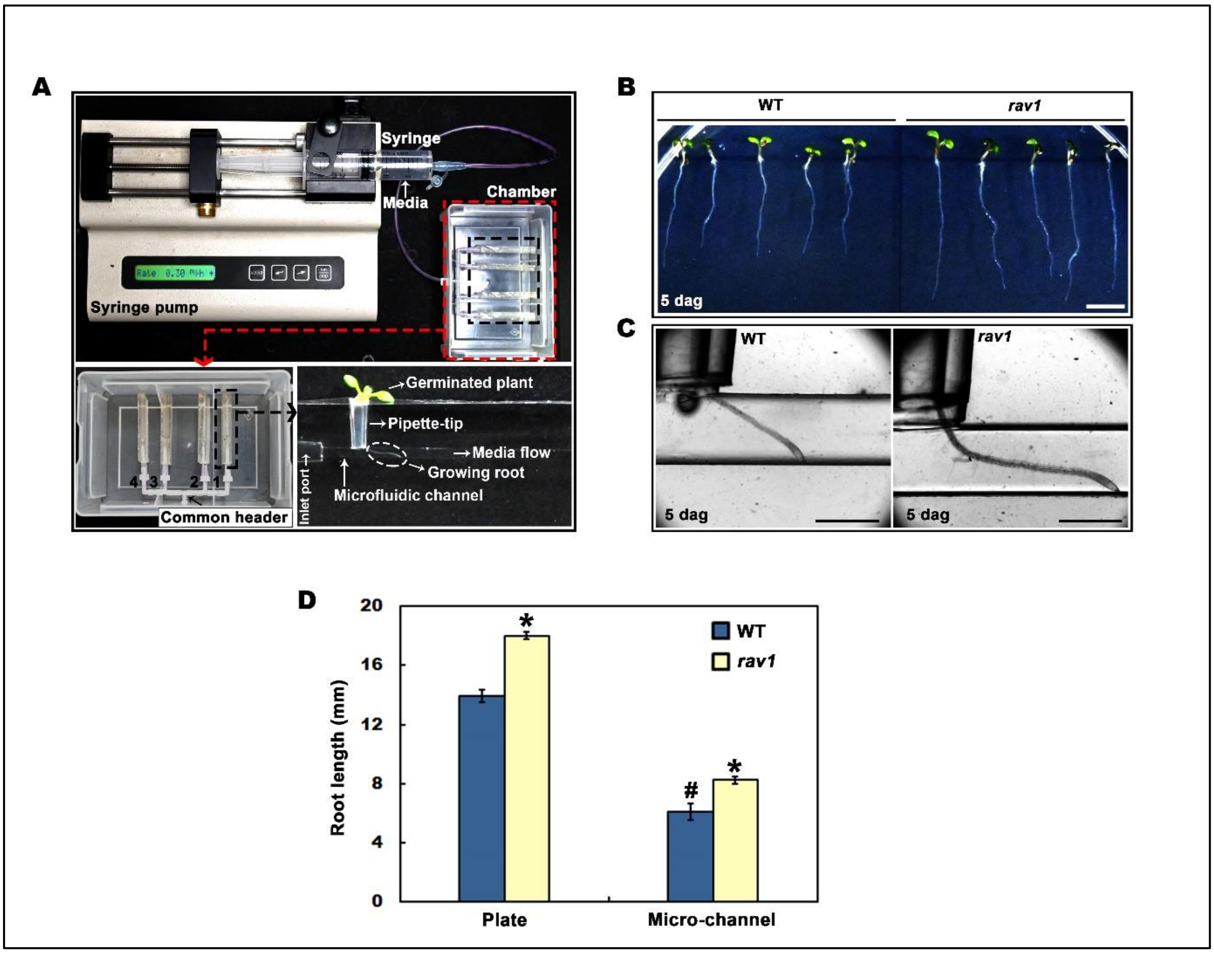


**Figure S2. Details of the microfluidic system and comparative root growth study. A.** Experimental set-up of Plant Root Microfluidic System (PRMS) attached to syringe pump. Four microfluidic channels in PRMS were connected to a common header through the inlet ports and the outlet ports were extended to the collection sump using connectors. This whole arrangement was placed in a closed chamber. Bottom left panel showing enlarged view of the chamber. Syringe pump containing 20 mL syringe was used to supply the ½-strength MS media solution in the channels through the common header. The media flow rate was set as 0.8 ml/hr from the pump, such that each of the four channels gets a flow rate of 0.2 ml/hr. Lower right panel shows enlarged view of a microfluidic channel with germinating plant and root growing into the channel. **B.** Primary root growth of WT and *rav1*, 5 dag. Surface sterilized Arabidopsis seeds were grown in vertical plates on solid MS medium supplemented with 3 % sucrose. 3 dag seedlings were transferred to solid ½-strength MS media plates and imaged after 2 days. Scale bars = 1 cm. **C.** WT and *rav1* roots growing into the microfluidics channel, 5 dag. Surface sterilized Arabidopsis seeds were germinated in 5 mm long pipette tip cone embedded in solid MS medium (3 % sucrose). 3 dag the pipette tip cone containing seedlings were inserted into microfluidic channel and supplied with ½-strength MS media for 2 days. Scale bars = 1 mm. **D.** Primary root length measurements of 5-day old WT and *rav1* grown either vertically on plate or on microfluidics channel was done and graphically plotted. The data represent mean of three biological replicate experiments on three separate occasions (n > 20) with standard error bars. Anova two-factor with replication method was used to calculate the variance between wild type and *rav1*. Variance in between WT and *rav1* is denoted by * and variance in between WT samples is denoted by # on corresponding graphs (P < 0.005).

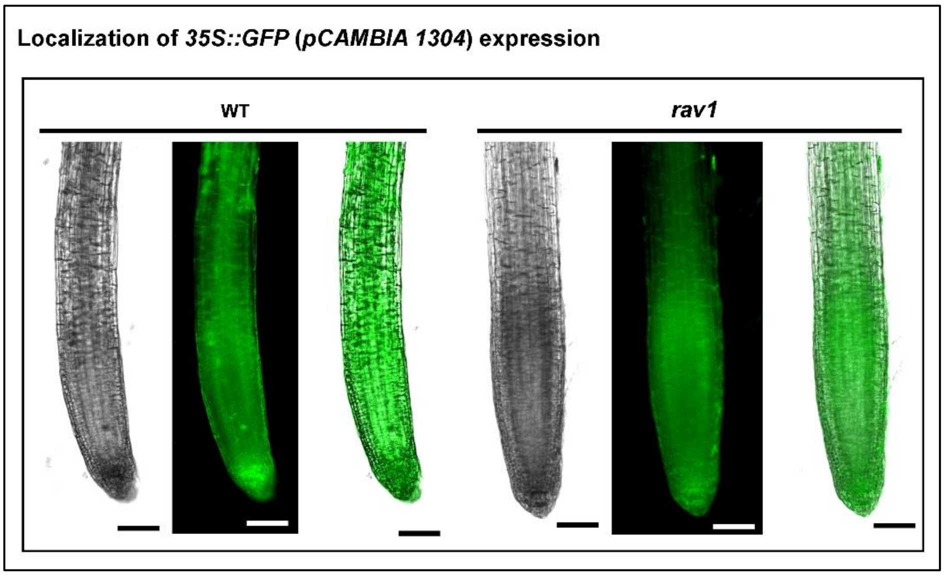


**Figure S3.** Expression of 35S::GFP (pCAMBIA1304) expression in wild type and *rav1* roots using hairy root transformation. Scale bars = 100 µm.

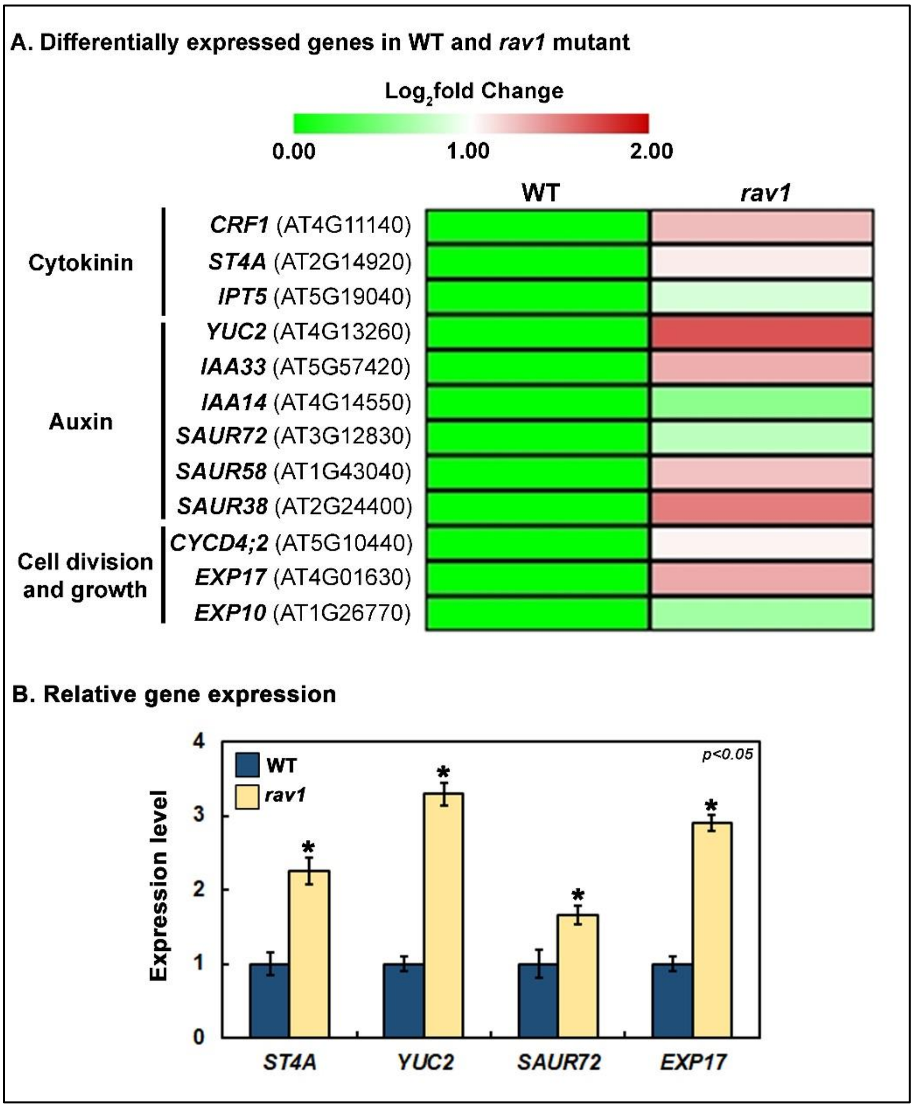


**Figure S4.** Comparative gene expression analyses from wild type and *rav1* roots 14 dag, involved in cytokinin and auxin hormonal pathways and related to cell division and growth. **A**. Heat map generated (MEV 4.6.0. software) from RNA-seq analyses, representing $\log_2$ fold change of transcripts (*rav1*/wild type). **B.** Validation of RNA seq data using qRT-PCR. The data represented are mean of three independent biological repeats with standard error bars. Statistical significance tested using paired two-tailed student's t-test and results with $P \leq 0.05$ are marked by *.

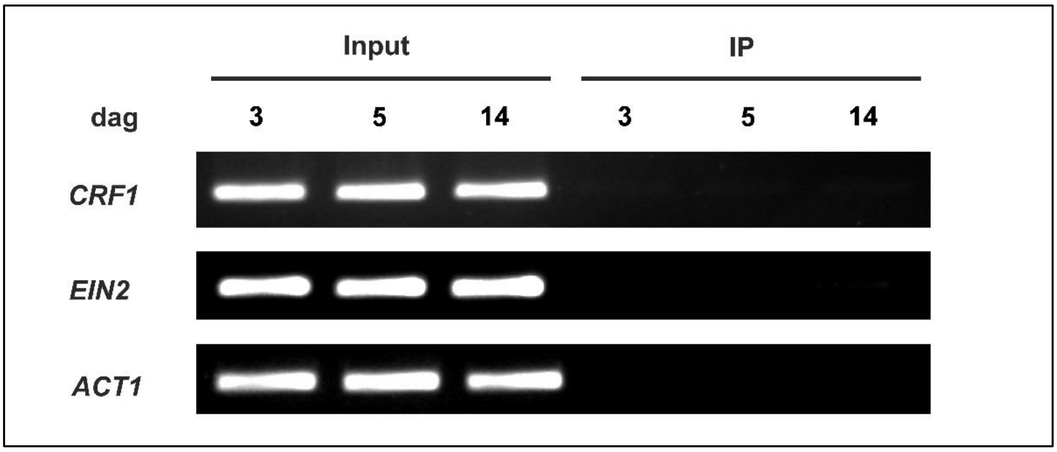


**Figure S5: ChIP-PCR for negative controls of RAV1 occupancy.** ChIP assay done using anti-RAV1 antibody followed by PCR at the *CRF1* locus using primers designed for regions that did not contain potential RAV1 binding sites. PCR was also done with a region of *EIN2* and housekeeping gene *ACT1* that do not have any known RAV1 binding sites. Panels show agarose gel images of PCR products for each gene done using input and immunoprecipitated DNA (IP) of wild type roots at 3, 5 and 14 dag.

**Supplemental Table S1: List of primers**

**A. For gene expression analysis:**

| **Accession number** | **Primer name** | **Sequence** |
|---|---|---|
| AT1G13260 | RAV1 FP | GGTCAGGATCAACAGTTGTAC |
| | RAV1 RP | CTTGCTACACACCAACGA |
| AT3G63110 | IPT3 FP | GAGTTTGACAGGTTTTTCAGGAAC |
| | IPT3 RP | AGAAACCGCGACACAGTATCTG |
| AT5G19040 | IPT5 FP | ACGGAGAGACTTCTTGAAACGG |
| | IPT5 RP | GGTCATCGCTGTAACTAAGGAAC |
| AT3G23630 | IPT7 FP | GAGCTCCATGAATATTTACGTAAC |
| | IPT7 RP | GACGATTCTCTCGCTTGGTCTC |
| AT3G16857 | ARR1 FP | CTACTCTTCTTCGTCCATGTCAAG |
| | ARR1 RP | AATCCCTTCCTGCTTAAGAAGTG |
| AT2G25180 | ARR12 FP | GCCAATACATTGAGTTCTCCAGCC |
| | ARR12 RP | TCTGTTCCTGCTTCATCGTGGAG |
| AT5G62920 | ARR6 FP | GAAGAAGATCAAAGAATCCTCAGC |
| | ARR6 RP | GAGTGAACAGGGTAGACATTCTC |
| AT1G04240 | SHY2 FP | ATGAGGGTCAAGGAATCTATGTG |
| | SHY2 RP | ACATATGAACATCTCCCATGGAAC |
| AT1G73590 | PIN1 FP | GGCGTTAAACCCAAGAATAATAG |
| | PIN1 RP | AAATATCACCGCAGTGCTAAG |
| AT4G37490 | CYCB1;1 FP | CAGACGAAAAGATGGAGAATATGG |
| | CYCB1;1 RP | CTTTGCACAGTCCATGAGCTGAG |
| AT4G11140 | CRF1 FP | GTTTCGGATTTTATCATTGGCGG |
| | CRF1 RP | CGCCGAAACTTTCACTCATCG |
| AT1G70940 | PIN3 FP | TCATTCAACAATCTATCTCCATTC |
| | PIN3 RP | GTAATGCGGCCTGAACTATAG |
| AT1G23080 | PIN7 FP | GGCAATGCCTAAAATTATTCAAC |
| | PIN7 RP | GCAATGCAGCTTGAACAATG |
| AT2G14920 | ST4A FP | CAGTTTGGTATTTTAGGCAAAGC |
| | ST4A RP | CAGATCCGCTATCTTCCTCTTCC |
| AT4G13260 | YUC2 FP | TCGTGACTCGGTACACGTATTAC |
| | YUC2 RP | CGGATACACCTTGATGTGTCCAC |
| AT4G01630 | EXP17 FP | ACTATCACAGCTACAAACTTCTG |
| | EXP17 RP | ATCCAAACTTTAGAGATCTCACC |
| AT3G12830 | SAUR72 FP | TCTCCTCCGTTCCGATTCACAGC |
| | SAUR72 RP | CCATGATTCGTTCGAAGACAAGG |
| AT1G16300 | GAPDH FP | AACAGGCGCTGCAAAGGCTG |
| | GAPDH RP | CGCCATGTGTTCTATCAGGTC |

**B. For ChIP assay:**

| Primer name | Sequence |
|---|---|
| CRF1 Region I FP | CACTTCACATATATTACTGTGTTCTG |
| CRF1 Region I RP | TTTGGAGTATTGATTAAATTTCTCTTG |
| CRF1 Region II FP | CGTGAAGGTGATGAATTACAAAG |
| CRF1 Region II RP | GGTATGGATGTTATATTCATAGTTAGC |
| CRF1 CNC FP | ACTCTTCTTCTTCTTCCTCTCTGTG |
| CRF1 CNC RP | ATCTGAGAACCTTGGAGACTCTAATG |
| EIN2 CNC FP | TTGCTCCTGCGCTTTATT |
| EIN2 CNC RP | CTGCTCCCAAATACCATCTC |
| ACT1 CNC FP | CTCCTCCTCCCTTCTTCTT |
| ACT1 CNC RP | TAGCTCAAGACACCCAGAT |

**C. For mutant line screening:**

| Primer name | Sequence |
|---|---|
| RAV1 gene-specific FP | AGCTTCCGTCGTCAAAATAC |
| N655012 T DNA RP | GGAAGCTAAACCGTTTGGTTATAC |
| ARR1-5’H2 FP | CTTCAAGCACTAGCCGTCACAGGTCAGT |
| ARR1-3’H3 RP | AATGTTATCGATGGAGTATGCGTCAAAGT |
| ARR1-3 RP | AGTGAAACCCGCTTCGGAGCTGTTAAC |
| CRF1-068G09 FP | TCCTTACGCAACAGATTCGTC |
| CRF1-068G09 RP | CGTGAGACAACACGTGATACG |

**D. For cloning:**

| Primer name | Sequence |
|---|---|
| ARR1 (DBD & TAD) FP | CACCATGAATTTGAAGAAACCGCGTG |
| ARR1 (DBD & TAD) RP | TCAAACCGGAATGTTATCGATG |
| RAV1 CDS FP | CAC CATGGA ATCGAGTAGC |
| RAV1 CDS RP | TTACGAGGCGTGAAAGATG |
| proRAV1-0.6 kb FP | CACCTAAGCATCAAAACTCAGGGG |
| proRAV1-0.8 kb RP | CACCGAGTAAAACACACACACTGAAGG |
| proRAV1 RP | GTAAGAATGCGTGTTTGAGG |
| proCRF1-1.2 kb FP | CGCGGATCCAGTGATTAGATATAAGTATATAAC |
| proCRF1-1.2 kb RP | CATGCCATGGGTGTGTTCTTCTTTGTGTCGTGTG |
| proPIN1 FP | CGGGATCCCTGTTGATGGAATATTGTGTTTC |
| proPIN1 RP | CATGCCATGGGAGAGAAGAGCTTTTGTCTTTAG |
